# FOXA1 mutations co-opt nascent transcription factor networks in partnership with androgen receptor to enhance prostate tumorigenicity

**DOI:** 10.1101/2025.11.13.688263

**Authors:** Erik M. Ladewig, Abbas Nazir, Tyler Park, Vinson B. Fan, Zhendong Cao, Jacob Hawk, Lauren Kelly, Robert Tjian, Christina S. Leslie, Charles L. Sawyers

## Abstract

Mutations in the pioneer transcription factor FOXA1, found in 10-40% of human prostate cancers, alter global chromatin accessibility and promote growth in prostate cells. Through analysis of a novel cohort of 874 primary and metastatic prostate tumors with somatic FOXA1 mutations, we confirm the high frequency of missense mutations (n=339) and indels (n=335) in the Wing2 region of the Forkhead domain, as well as frameshift mutations that truncate the C-terminus (n=287). To investigate the transcriptomic consequences of each mutation subgroup as well as elevated levels of wild-type FOXA1 (WT) (a fourth well documented subgroup), we performed single nucleus multiome sequencing in primary mouse organoids following inducible expression of representative alleles. Whereas each mutant induced distinct transcriptomic and DNA accessibility features, a prominent feature of all mutants was perturbed epithelial lineage specification, ranging from basal-like fates in cells expressing indel mutants to secretory (L1-like) luminal fates in cells expressing C-terminal truncation, missense mutations or excess WT. Integrated RNA-seq, ATAC-seq and ChIP-seq analysis of L1-like fate specification revealed enrichment of a composite androgen receptor-FOXA1 hybrid motif and cooperativity with the Oct family transcription factor POU2F1. Importantly, L1-like fate specification is seen *in vivo* tumorigenicity assays where, in combination with *Trp53*/*Pten* loss, expression of these mutants results in a histologic switch from basal-like to secretory luminal histology.

## Introduction

FOXA1 encodes a developmentally critical pioneer transcription factor (TF) that becomes an oncogenic driver in human prostate and breast cancer through distinct types of genomic alterations (*1–3*). FOXA1 mutations are present in ∼12% of prostate cancer patients in Western cohorts and ∼40% in Asian cohorts. Curiously, this ethnicity difference is also seen with ERG translocations but in the reverse direction, common in Western cohorts but rarer in Asian cohorts. Notably, FOXA1 and ERG alterations are mutually exclusive across ethnicities and collectively account for ∼60% of all prostate cancer worldwide.

The most frequent FOXA1 alterations are missense or indel mutations in the Wing2 region of the Forkhead domain (*1*, *4*). Nonsense mutations that lead to C-terminal truncation define a third group, which is reported to be exclusive to metastatic disease (*2*, *5*). A fourth group of FOXA1 alterations is defined by elevated levels of WT FOXA1 mRNA and protein - due to focal gene amplification, FOXA1 promoter or 5’ UTR mutations, or translocations.

Several groups have explored the biological consequences of mutant FOXA1 expression in cell lines, organoids and mice. We previously used primary mouse prostate organoids to show that mutant FOXA1 expression results in widespread chromatin accessibility changes enriched for canonical and non-canonical Foxa1 binding motifs within days of induction, leading us to postulate that sequence-specific DNA binding of mutant FOXA1 is a key first step for aberrant pioneering (*1*). We and others have also reported mutant-specific effects on prostate epithelial cell fate, ranging from activation of basal-like to pro-luminal gene expression programs depending on the mutation, as well as tumorigenicity in mice (*1*, *2*, *5*, *6*).

Here we extend these analyses by first reporting on a larger cohort of patients with FOXA1 mutant prostate cancers (n=874) compared to earlier reports with cohorts ranging from 89 to 370 FOXA1 mutant cases (*1*, *2*, *4*). The new analysis confirms the high frequency of missense, indel (in frame), and frameshift mutations reported previously but now with greater precision. One insight is the fact that nearly all indels map to either M253 or F254 (n=335), underscoring the importance of these two residues in Wing2 function. A second new insight is the presence of truncating (nonsense) mutations at equal frequency in primary cancer and metastasis, in contrast to earlier reports (*2*, *5*).

To gain novel mechanistic insight, we performed systematic transcriptional and TF motif accessibility analyses at single-cell resolution (scRNA+ATAC-seq) because this technology can reveal underlying cell state transitions. This integrated single-cell approach revealed that all FOXA1 mutants have a profound impact on prostate epithelial lineage fate. The most consistent and prominent lineage effects were rapid changes in the percentage of basal cells (increased by expression of mutants at M253 and E255, representing the two indels) or L1-like secretory luminal cells (increased by expression of mutants with C-terminal truncation, H247 missense or excess WT). The appearance of L1 cells in organoid culture is remarkable because, although L1 is the predominant cell type in mouse and human prostate glands *in vivo*, they are poorly represented in organoid culture, likely due to growth conditions that favor self-renewal of stem-like (L2) cells.

Through integrated RNA-seq, ATAC-seq and ChIP-seq analysis, we identified a non-canonical composite FOXA1-AR binding motif, and the homeodomain POU2F1 TF as uniquely associated with and required for induction of L1 luminal fate. Direct comparison of the truncation mutant (G275X) with WT revealed that absence of the C-terminus shortened the time required for cells to reach the L1 state (24 hours versus 5 days) as well as chromatin residence time in single molecule tracking (SMT) experiments, consistent with a “fast” mechanism postulated earlier (*2*). These mutants also induce an L1-like state *in vivo*, initiating a phenotypic switch to secretory luminal histology in a background of *Pten*/*Trp53* loss, a widely used model of prostate cancer with predominantly basal-like features. Our analyses therefore uncovered a novel transcription factor network comprising FOXA1-AR and POU2F1 that both drives the L1 luminal secretory fate and enhances tumorigenicity downstream of mutant FOXA1 expression.

## Results

### Updated map of FOXA1 mutants in human prostate cancer

Prior work mapping the location of FOXA1 mutations in human prostate cancer relied on cohorts with ∼89-370 independent FOXA1 mutant cancers, including primary tumors and metastases (*1*, *2*, *4*). Leveraging the paired germline and tumor sequencing routinely performed at Memorial Sloan Kettering Cancer (MSK) using the MSK-IMPACT test (*7*), we identified 686 primary prostate cancers and 305 metastatic prostate cancers with somatic FOXA1 mutations (**Fig 1A-B, Table S1, Methods**). Consistent with earlier analyses of smaller cohorts, mutations fall into three subgroups: missense (n=339), indels (inframe) (n=335) and truncation (nonsense) (n=287). As expected, missense mutants and indels, which collectively account for 68% of FOXA1 mutations in this cohort, map almost exclusively to Wing2 in the Forkhead domain, whereas truncation mutations (29% of FOXA1 mutations) are spread across the C-terminus distal to Fkhd. Of note, truncation mutants are present with equal frequency in primary tumors and metastases, a finding that differs from another cohort (*2*, *5*). Another striking finding is that nearly all indels map to just two residues within Wing2 (M253, E255). A cohort of this size linked to clinical outcome data provided an opportunity to explore potential prognostic differences across mutation subgroups. Toward that end, we found that patients with tumors with indels (inframe mutations at M253 or E255) have improved survival compared to those with truncating (nonsense) mutations (**Fig S1A**).

**Fig. 1.**
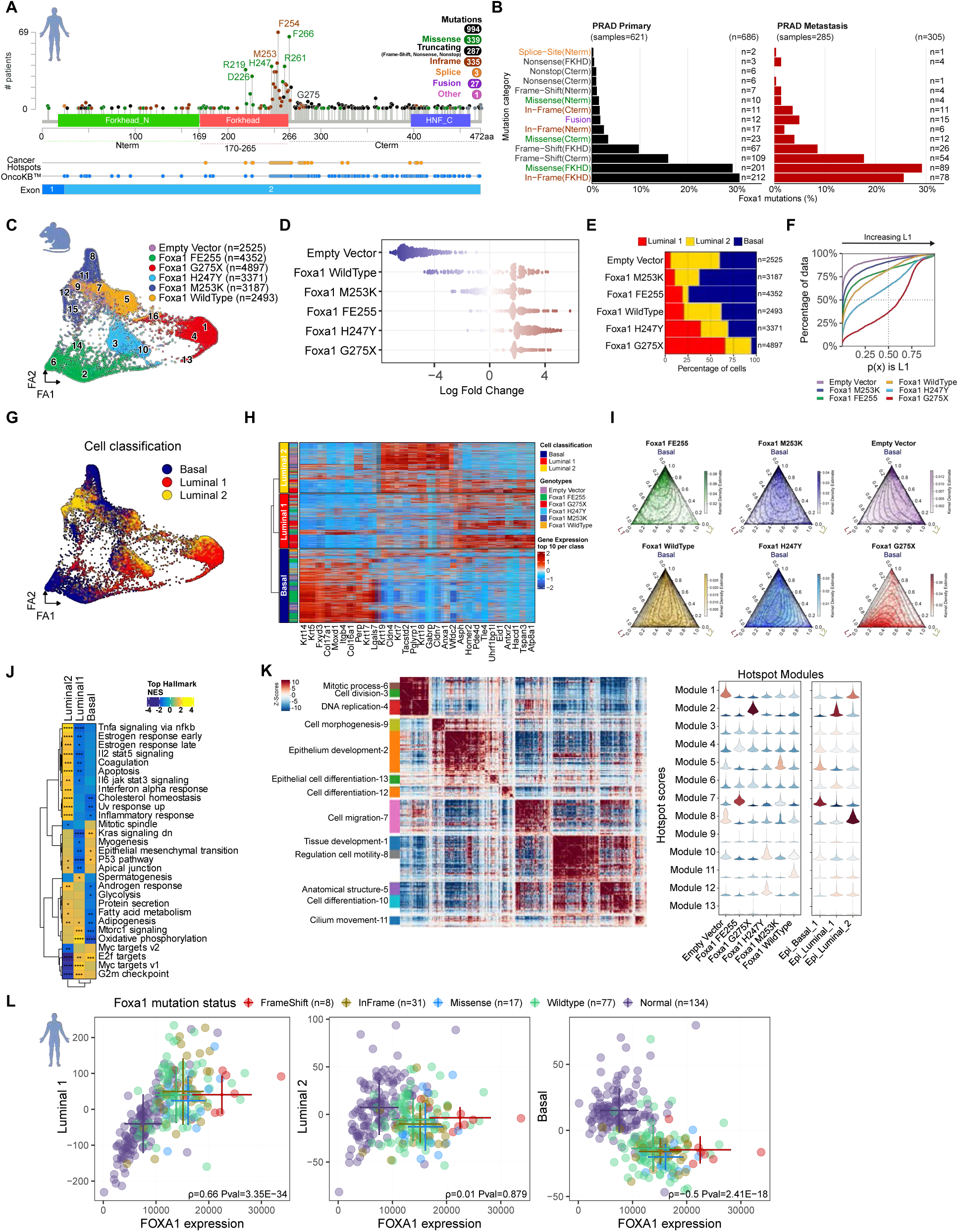
FOXA1 mutations drive epithelial cell specific transcriptomes in prostate organoids. (A) A lollipop plot depicting a simplified view of human FOXA1 gene, its protein domains, and the location and number of mutations seen in prostate cancer cohorts obtained through cbioportal.org (see Methods). Frequent mutations in this study are labeled. (B) Barplots categorized by mutation type in either primary or metastatic prostate adenocarcinoma (PRAD) ordered from least (top) to most frequent (bottom). (**C**) Force directed layout of scRNA-seq of mouse prostate derived organoid cells colored by genotype. Labels from Leiden clustering are overlaid. (**D**) Cell neighborhood analysis graph produced by Milo showing neighborhood log fold changes grouped by genotype (*35*). (**E**) Bar plot of cell type proportions per genotype ordered by increasing L1 ratio. (**F**) ECDF of genotype probabilities to be L1 luminal cell. (**G**) Force directed layout of scRNA-seq colored by basal, L1, or L2 cell types. (**H**) Heatmap of most significantly differential genes from L1, L2, and basal grouped by genotype. (**I**) Ternary plots depict a point for every cell that represents a probability of the cell being basal, L1, and L2 on an equilateral triangle. Density of cell type probabilities are shown in contours and color concentration. (**J**) GSEA categories shown on heatmap indicating enrichment in L2, L1, and basal cells. (**K**) Heatmap of hotspot gene modules detected and annotated with significant Gene Ontology terms (left panel). Violin plots separated by hotspot modules with hotspot scores on y-axis and sample genotypes on x-axis (right panel). (**L**) Human prostate cohort scatterplots showing the distribution of individual samples stratified by *Foxa1* mutation categories: Frameshift (red), In-Frame (orange), Missense (blue), Wildtype (green), and Normal (purple). Except for normal tissue, all categories were defined using prostate cancer (PCa) samples, including "wildtype" which refers to FOXA1-intact. Error bars represent the mean ± standard deviation for each group. *FOXA1* expression correlates with L1 activity in both normal and tumor samples, with tumors showing elevated levels. Frameshift mutants display the highest *FOXA1* expression, suggesting enhanced L1 activation. L2 scores correlate with *FOXA1* in normal tissue but not tumors, while basal activity is largely uncoupled and lower in tumors.

### Oncogenic FOXA1 mutants promote a spectrum of luminal and basal lineage fates

Building on our earlier work documenting gain of function in chromatin opening activity of FOXA1 mutations found in prostate cancer patients, we selected a representative Wing2 missense mutation (H247Y), the two indel mutations (M253K, FE255), a truncation mutation just downstream of the Forkhead domain (G275X) and WT for deeper investigation into how these chromatin accessibility changes impact transcription and oncogenesis. Because of their presumed role in disease initiation, we examined their activity in freshly isolated primary mouse prostate cells grown in organoid culture, a system that we and others have recently optimized to study dynamic events in prostate cancer initiation and progression *in vitro* and *in vivo* following orthotopic transplantation (*8*), (*9*), (*10*). Robust expression of Flag-tagged WT and each mutant allele and confirmed 5 days after doxycycline induction at levels well above endogenous FOXA1, albeit with WT somewhat lower than each of the mutant subclasses (**Fig S1B-C**).

We next used single-cell multiome sequencing to simultaneously profile the transcriptome and chromatin accessibility differences of each in comparison to an empty vector (EV) control. We first clustered cells by their single-cell transcriptomes and visualized using a 2D force-directed layout (with ForceAtlas, FA) (**Fig 1C**). Cells clustered primarily by mutant, exemplified by the fact that 12/16 (75%) clusters contained 75% or more cells from a single genotype (**Fig S1E-F**). Indeed, only two clusters – cluster 14, enriched in cell cycle genes, and cluster 16, dominated by ribosomal RNA – contained cells from all samples (**Fig S1D,1G**). To better compare the differences between mutant and EV control cells, we performed neighborhood-based differential abundance tests for cells of each mutant genotype versus EV using MILO (see **Methods**). The computed log fold changes over neighborhoods show cells expressing FE255, H247Y and G275X have diverged the most from EV control (**Fig. 1D**).

In searching for genes that may account for this mutant-specific clustering, we noted the most statistically significant differentially expressed genes (DEG) that distinguish the FOXA1 clusters included luminal (*Krt8*, *Krt19*) and basal cell (*Krt5*, *Krt14*, *Krt15*) marker genes, suggesting that different FOXA1 mutations bias cells to distinct epithelial fates (**Fig. S1H**). Prostate tissue contains two major epithelial cell types: (i) basal cells, which line the basement membrane, and (ii) luminal cells which secrete the protein constituents of seminal fluid. The luminal cell compartment includes two primary subtypes: (i) differentiated secretory cells (called L1 or distal Lum), which account for >90% of luminal cells in the prostates of hormonally intact males and (ii) stem-like cells (called L2 or proximal Lum) (5-10% of luminal cells) (*11*), (*12*). Importantly, the ratio of L1 to L2 cells seen *in vivo* is reversed in organoid culture, where L1 cells are poorly maintained due to growth conditions that favor growth of basal and L2 luminal cells (*13*). Indeed, we found that L1 cells (annotated using literature-derived signature for L1, L2 and basal cells) account for only 6% of epithelial cells in EV cultures. Conversely, cultures expressing each of the five mutant alleles (including elevated WT) had increased levels of L1 cells, ranging from 10% (M253K), 19% (FE255), 20% (WT), 39% (H247K) and 66% (G275X) (**Fig. 1E**). These findings were supported by a Gaussian mixture model, using the ECDF of the L1 posterior probability (**Fig 1F, S1I–K, Methods**), and further validated by morphological evidence of large cystic lumens in organoid bright-field images (**Fig S1L**), consistent with enhanced secretory activity. UMAP visualization revealed cell type clusters were largely driven by epithelial lineage phenotypes suggesting FOXA1 mutant transcriptomes generate variation among defined cell types (**Fig 1G**). Differential expression analysis of basal, luminal 1 (L1), and luminal 2 (L2) cell types identified basal (Krt5, Krt14) and luminal genes (Krt18, Krt19) and L1 (Tspan3, ATp8a1) and L2 specific (Cldn4, Tacstd2) genes (**Fig 1H, Table S2, Methods**). Ternary plots further quantified these identities, showing that WT, H247Y, and G275X cells all clustered preferentially toward the L1 vertex (**Fig 1I**). While it is possible that the expanded proportion of L1 cells is due to faster proliferation, the magnitude and speed with which this occurs (11-fold over 5 days for G275X) coupled with the amount of chromatin remodeling supports a lineage conversion model.

Although an increase in L1 cells were seen across all mutants, the FE255 and M253K alleles also led to a significant increase in basal cells (**Fig 1E**), similar to results reported independently with a F254E255 mutant (*6*). Thus, a primary consequence of oncogenic FOXA1 expression is a shift in epithelial lineage specification, with indel mutants (M253K, FE255) showing basal skewing whereas truncation (G275X), missense (H247Y) and WT all display secretory L1-like skewing.

To move beyond lineage specification, we performed Gene Set Enrichment Analysis (GSEA) and found upregulated biological pathways such as oxidative phosphorylation, glycolysis, androgen response and mTORC1 signaling in the L1 cell clusters from G275X and H247Y (clusters 4 and 10 respectively) as well as MYC targets and E2F targets in all clusters of FE255 and G275X (**Fig 1J-K, S1M**). We then used our single-cell data to see whether shared and distinct gene expression programs are organized into biologically meaningful modules across Foxa1 mutants. Hotspot analysis (*14*) yielded 13 modules shared across all mutants that are associated with oncogenesis, including mitotic processes (module 6), cell division (module 3) and DNA replication (module 4) (**Fig. 1K, left panel**). Other programs such as epithelium development (module 2) and cell migration (module 7) were linked to specific mutants (G275X and FE255, respectively). Interestingly, these mutant-specific modules are similarly enriched in the epithelial lineage primarily associated with each of those mutants (L1 cells and basal cells respectively) (**Fig. 1K, right panel**).

To contextualize our murine findings within human disease, we analyzed data from the Chinese Prostate Genome and Epigenome Atlas (*4*), the largest cohort to date with matched tumor/normal expression profiles and annotated FOXA1 mutation status with sufficient numbers of FOXA1 mutant cases (56 FOXA1-mutant tumors from a RNA-seq cohort of 134 tumor/normal pairs, see Methods) to make meaningful comparisons across mutation subgroups. We evaluated FOXA1 expression and L1, L2, and basal epithelial gene signatures across distinct mutation categories (**Fig. 1L, Table S3, Methods**). Except for normal tissue, all categories represent prostate cancer (PCa); thus, "Wildtype" refers to FOXA1-intact PCa. L1 activity positively correlated with *FOXA1* expression in normal and in tumor samples. Notably, tumors with FOXA1 frameshift mutations showed the highest FOXA1 expression as well as L1 activation. L2 signature scores correlated with *FOXA1* in normal tissues but not in tumors. Basal signature activity was inversely associated with FOXA1 expression, which was generally reduced in tumor tissue. Unlike mouse organoids, patients with inframe mutants did not have evidence of basal lineage skewing, albeit with a limited number of cases.

In summary, all subclasses of FOXA1 mutants represented by these alleles activate transcriptomic changes that skew the relative proportion of basal, progenitor-like (L2) luminal and differentiated/secretory (L1) luminal cells in primary organoid culture, together with induction of mitotic programs associated with oncogenesis. Notably, the L1 skewing profile matches transcriptomic features in FOXA1-mutant human cancers.

### Chromatin opening initiated by mutant FOXA1 alleles reveals lineage specific TF motifs

Having characterized the transcriptional changes induced by each FOXA1 allele, we next analyzed chromatin accessibility from the organoid multiome dataset. Cells were subjected to quality filters (**Fig. S2A**), represented by variable genomic tiles, clustered using a graph-based approach, and visualized by Uniform Manifold Approximation and Projection (UMAP) (**Fig. 2A, Fig. S2B**). This representation primarily clustered cells by genotype, revealing distinct chromatin accessibility profiles among mutants, consistent with the large number of mutant-specific accessible peaks detected in the scATAC-seq data (**Fig. S2C**). The FOXA1 G275X mutant exhibited the highest number of differential peaks relative to the Empty Vector (EV) control, and the largest cluster (C1) that overlapped with a dense region of L1 cells (**Fig 2B, Fig. S2D-E**). Overall, G275X showed the most extensive mutant specific accessibility changes in the scATAC-seq peak atlas. The presence of coordinated accessibility and transcriptional shifts across nearly all cells within each genotype suggests that FOXA1 pioneering activity operates uniformly within individual cultures.

**Fig. 2.**
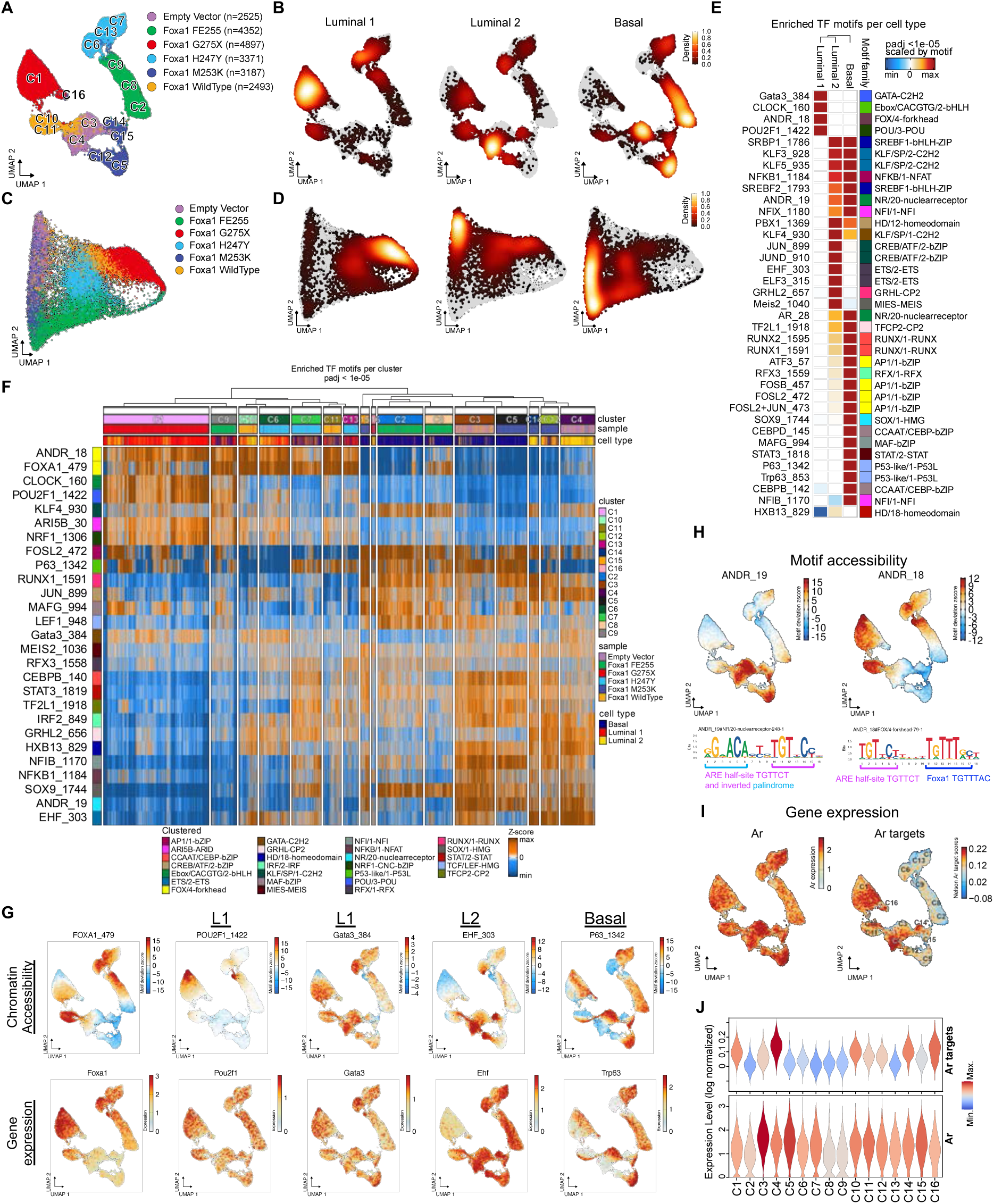
FOXA1 single-cell chromatin accessibility reveals distinct transcription factor motifs that integrate luminal / basal expression programs. (**A**) scATAC-seq uniform manifold approximation and projection (UMAP) of mouse prostate organoids (**B**) UMAPs colored to show density of cells in each epithelial subtype. (**C**) scATAC-seq force directed layout projected from alternative CellSpace embedding (**D**) Force-directed layout generated from CellSpace cell density colored for each epithelial subtype. (**E**) Heatmap of significant motifs assessed by epithelial subtype and colored according to -log10(adjusted p-value). Motifs are also labeled by motif families. (**F**) Average TF motif accessibility of top motifs that are both significantly accessible and correlated with expression per genotype. (**G**) Individual UMAPs of chromatin accessibility (top) and gene expression (bottom) for several motifs significantly accessible according to cell type. (**H**) AR Canonical motif ANDR_19 and composite motif ANDR_18 accessibility assessed by chromVAR each shown on a UMAP (top left, top right, respectively). The ANDR_19 motif sequence with bracketed androgen response element (ARE) half-site and inverted palindromic ARE site (bottom left). ANDR_18 hybrid motif of ARE half-site and Foxa1 (bottom right). (**I**) scATAC-seq UMAPs showing *Ar* expression, indicating broad expression in genotypes (left). scATAC-seq UMAPs showing high AR target gene expression within regions containing mostly L1 and L2 cell types (right). (**J**) Violin plots of *Ar* expression (left) and AR target expression (right).

The discovery that FOXA1 mutants with similar transcriptomic programs (e.g., L1 luminal with G275X, H247Y and WT) exhibit distinct ATAC profiles led us to investigate whether TF motifs in newly accessible regions drive FOXA1-dependent expression changes. Using the sequence-informed scATAC-seq embedding algorithm CellSpace (*15*), which maps DNA k-mers and cells into a shared latent space, we uncovered striking chromatin state overlaps among mutants. These states, organized into basal, L1 luminal, and L2 luminal clusters anchored at the three corners of the embedding, reveal that while each FOXA1 mutant exhibits unique chromatin accessibility, shared lineage-specific TF binding signals underlie the three canonical epithelial states (**Fig. 2C-D**).

To further investigate chromatin binding we identified key TF motifs with chromatin accessibility linked to TF expression within each epithelial state (Methods). Basal-like cells showed open chromatin with TRP63 (a basal cell marker) and SOX9 motifs (**Fig 2E-G)**. In Ar+ basal and L2 cells (C3, C4, C5, C14), we observed enrichment of the canonical AR motif ANDR_19 consisting of an inverted palindromic repeat followed by the androgen receptor element (ARE) half-site (AGAACA…TGTTCT). L2 cells also displayed motifs for ELF3 and EHF (ETS family TFs), paralleling luminal progenitor cells in breast tissue (*16*), along with HOXB13 (an AR cofactor and regulator of AR transcriptional activity (*17*), (*18*).

Surprisingly, in L1 cells, the ANDR_19 motif was inaccessible despite strong AR target gene activation (**Fig 2H**). Instead, these cells showed enrichment of an alternative hybrid Ar:Foxa1 motif (ANDR_18: TGTTCT…TGTTTG) (**Fig. 2E-F**). L1 cells (G275X cluster) exhibited the highest AR target gene expression, aligned with the accessibility of ANDR_18 (**Fig 2H**). In contrast, basal clusters (C3, C5; EV, M253K genotypes) enriched for ANDR_19 had lower AR target expression despite higher Ar mRNA levels (**Fig 2J**).

As suggested from the CellSpace analysis, we also noted lineage-specific enrichment of other TF motifs that include POU2F1/OCT1, a TF involved in regulation of development (*19*) and GATA3, a TF expressed in breast and prostate luminal cells (**Fig. 2E-G)**. Despite evidence implicating *Gata2* in AR-dependent prostate cancer (*20*, *21*), we did not find evidence of *Gata 2* expression in the mouse organoid models (**Fig S2F)**. In summary, lineage-specific enrichment of ANDR_18 versus ANDR_19 motifs coupled with different magnitudes of AR target gene activation (despite comparable levels of Ar expression) suggest that FOXA1 perturbs the AR cistrome to dictate distinct epithelial cell states in cooperation with additional partner TFs.

### FOXA1/AR colocalization at composite binding motifs is associated with lineage specific TF expression

Having identified a hybrid AR:FOXA1 motif linked to enhanced AR output and an L1 luminal cell fate, we next conducted chromatin immunoprecipitation sequencing (ChIP-seq) experiments to search for biochemical evidence of AR-FOXA1 cooperativity, focusing on FOXA1 mutants that promoted high L1 cell content (G275X, H247Y and WT), with M253K and EV serving as controls. Analysis of 67,811 AR ChIP-seq peaks grouped into ten clusters revealed that motifs enriched in L1 cells from scATAC-seq data (**Fig 2E**) were also detected in Ar ChIP-seq (**Fig S3A-B**). These include the composite hybrid motif ANDR_18, along with POU2F1 and GATA3 transcription factor (TF) motifs (clusters 2, 4, 5, 6) (**Fig 3A**). Interestingly, these same motifs were enriched in FOXA1 ChIP-seq data from G275X and WT organoids, strongly supporting co-binding by AR and FOXA1. In contrast, the canonical AR motif ANDR_19 was absent from these peaks but significantly enriched in EV and M253K organoids (cluster 7), where ANDR_18 was depleted. This divergence suggests distinct motif preferences between L1-promoting FOXA1 mutants and non-L1 states. To explore whether usage of ANDR_18 or ANDR_19 motifs is associated with differential response to AR inhibition, we performed proliferation assays in the presence or absence of enzalutamide. G275X and WT expressing organoids (both with ANDR_18 usage) were more sensitive to AR inhibition than EV or FE255 (ANDR_19) **(Fig S3F)**, consistent with the fact that L1 cells in the normal prostate display greater castration sensitivity *in vivo* (*11*). However, the sensitivity of H247Y organoids (also ANDR_18 and pro-L1), was indistinguishable from EV, suggesting greater complexity within L1-like signatures that is not always linked with AR dependence. Indeed, we previously showed that a subset of L1 cells in the normal mouse prostate persist following castration (*11*).

**Fig. 3.**
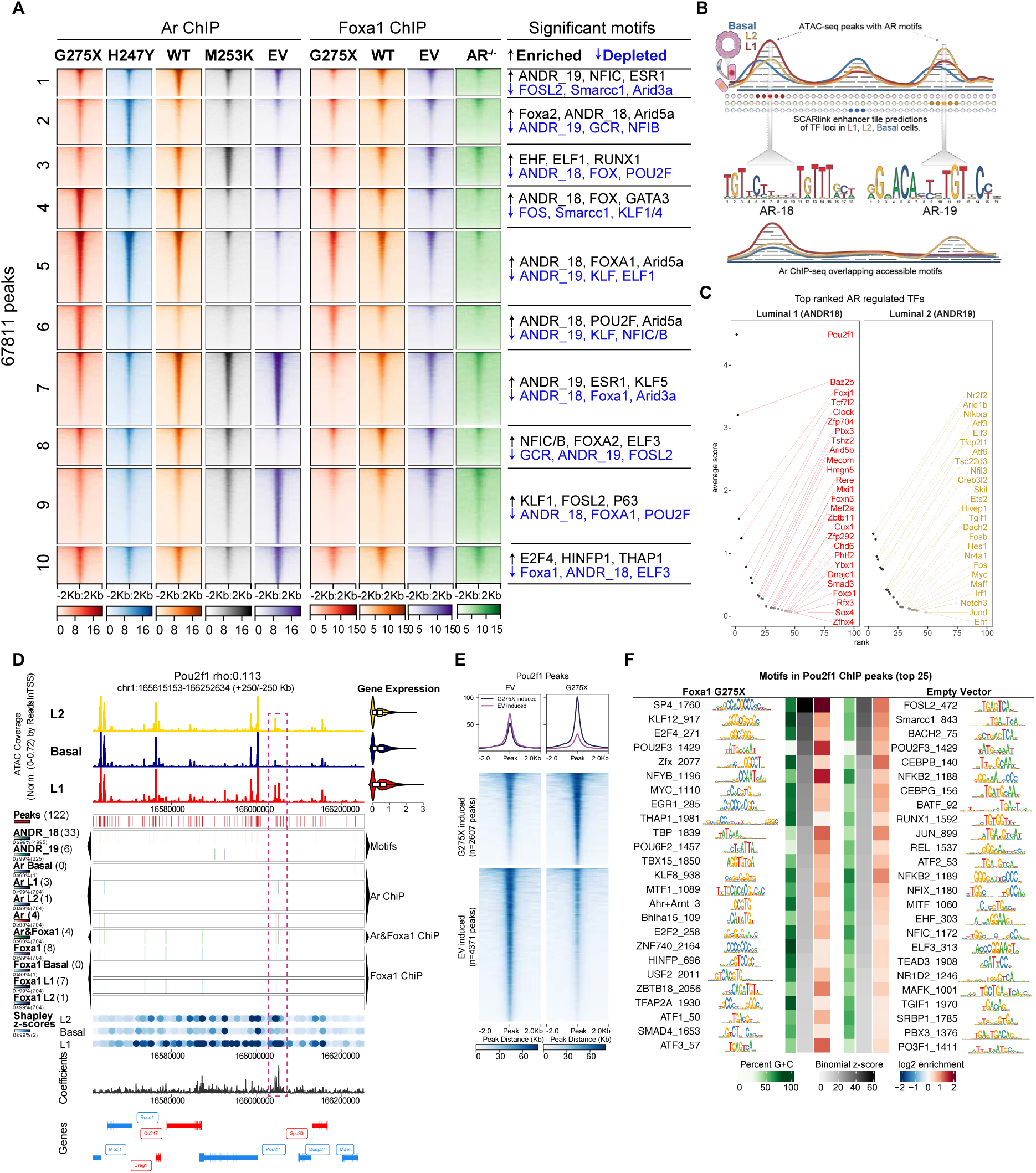
FOXA1 G275X cooperates with androgen receptor by enriching for an alternative Ar motif and reinforces Ar target gene expression. (**A**) Tornado plots of AR ChIP-seq (left) or FOXA1 ChIP-seq (right) data separate by AR driven clustering. Significantly accessible motifs detected in AR ChIP-seq data per cluster are shown on the far-right side and indicated as more binding (enriched) or less binding (depleted). (**B**) Schematic of scATAC-seq, ChIP-seq, and SCARlink analysis to nominate genes with Ar regulated enhancers. scRNA-seq gene expression is estimated from scATAC-seq for each cell type (L1, L2, basal). scATAC-seq is tiled into 500nt bins using SCARlink around gene bodies (+-250KB) where each tile is then scored using Shapley. Significant shapley scores designate candidate enhancers per gene that are overlapped with accessible AR motifs (ANDR18, ANDR19) and AR ChIP-seq peaks. The combined scores are averaged and used to rank potential Ar regulated TFs. (**C**) Average SCARlink scores per transcription factor that contain the ANDR_18 motif in AR ChIP peaks within all luminal 1 regions (left panel) or basal regions (right panel). Each transcription factor is ranked according to average scores (methods). (**D**) *Pou2f1* genomic locus (+/- 250 KB) with tracks listed in order from top to bottom: 1) normalized chromatin accessibility for scATAC-seq separated into L2, basal, and L1, 2) significant peak detection, 3) motif detection for ANDR_18, 4) ANDR_19, and 5) Foxa1. Then bulk AR ChIP-seq tracks separated into 6) basal, 7) L1, 8) L2, and 9) total AR, the 10) intersection of peaks in both AR and FOXA1, 11) ChIP-seq for total FOXA1, FOXA1 separated into 12) basal, 13) L1, 14) L2. Next are Shapley values from SCARlink loci separated to 15) L2, basal, and L1, then 16) SCARlink coefficients track and finally 17) gene annotations track. I) Top ranked AR linked genes in L1 luminal peaks utilizing hybrid motif ANDR_18 or canonical AR motif ANDR_19 are shown on left and right side, respectively. (**E**) Tornado plots showing peaks that are significantly differential (adjusted p-value < 0.05 & Log2FC > 0.5) in Empty Vector (EV, left) versus FOXA1 G275X (right) (**F**) POU2F1 ChIP motif enrichment performed in G275X and Empty Vector mouse prostate organoids (left, right respectively). Motifs shown are the 25 most highly significant as ranked by binomial z-score and ordered accordingly from greatest (top) to least (bottom) significant.

The selective enrichment of POU2F1 and GATA3 motifs in WT, H247Y, and G275X organoids raised the possibility that these TFs act as downstream targets of the hybrid ANDR_18 motif. To test this hypothesis, we conducted a genome-wide SCARlink analysis, integrating scATAC-seq and scRNA-seq data to identify lineage-specific enhancers in L1, L2, and basal states. Enhancers were segmented into 500-nt tiles and ranked using SCARlink-derived Shapley scores, then cross-referenced with Ar ChIP-seq peaks and accessible AR motifs (ANDR_18 or ANDR_19) to pinpoint Ar-regulated enhancers of TF genes (**Fig 3B**) (*22*); Methods). This analysis identified POU2F1 as the top AR-regulated TF candidate in L1 cells associated with the ANDR_18 composite motif (**Fig 3C-D**, left). Similarly, investigation of the ANDR_19 motif in L2 cells identified three ETS family genes (*Elf3*, *Ets2*, and *Ehf*) as potential AR-regulated targets (**Fig 3C**, **right**). ATAC-seq data confirmed selective enrichment of Elf3 and Ehf motifs in L2 cells (**Fig 2E**), highlighting their relevance to L2-specific gene regulation.

Further analysis underscored the potentially pivotal role of POU2F1 in L1 cells. For example, the GATA3 motif, prominently enriched in L1 cells (**Fig 2E**), was also present in L2 cells (**Fig 2F**), aligning with its established function in luminal cell identity in breast and prostate tissues (*23*), whereas the POU2F1 motif exhibited exclusive enrichment in L1 cells (**Fig 2E-G**, **Fig 3D**). Consistent with the motif analysis, POU2F1 ChIP-seq revealed changes in the genomic distribution of POU2F1 in G275X cells compared to EV as well as enrichment for novel motifs for multiple TFs including KLF (KLF12, KLF8), E2F (E2F4, E2F2), and MYC (**Fig 3E-F**). Of note, FOXA1 and POU2F1 motifs were also highly enriched in a bulk ATAC-seq study of 26 primary human prostate adenocarcinomas (100, 84 percentiles, resp.) (*24*). Together, these findings point to POU2F1 as a central AR-regulated TF in defining L1 identity.

### POU2F1 is a critical downstream effector of the pro-luminal (L1) phenotype

Having implicated *Pou2f1* as a direct AR-FOXA1 target gene, we next asked if POU2F1 is required for induction of the L1 transcriptional program through CRISPR targeting. We selected G275X for these studies based on robust induction of the L1 signature (**Fig 1E-F**). Although not directly implicated as an AR-FOXA1 target by SCARlink (**Fig 3C**), we also included *Gata3* because its motif was also selectively enriched in L1 cells (**Fig 2E**). After confirming CRISPR deletion of each TF at the protein level (**Fig S4A**), we performed single-cell multiome (scRNA+scATAC) sequencing 5 days after Dox-induction (**Fig 4A**).

**Fig. 4.**
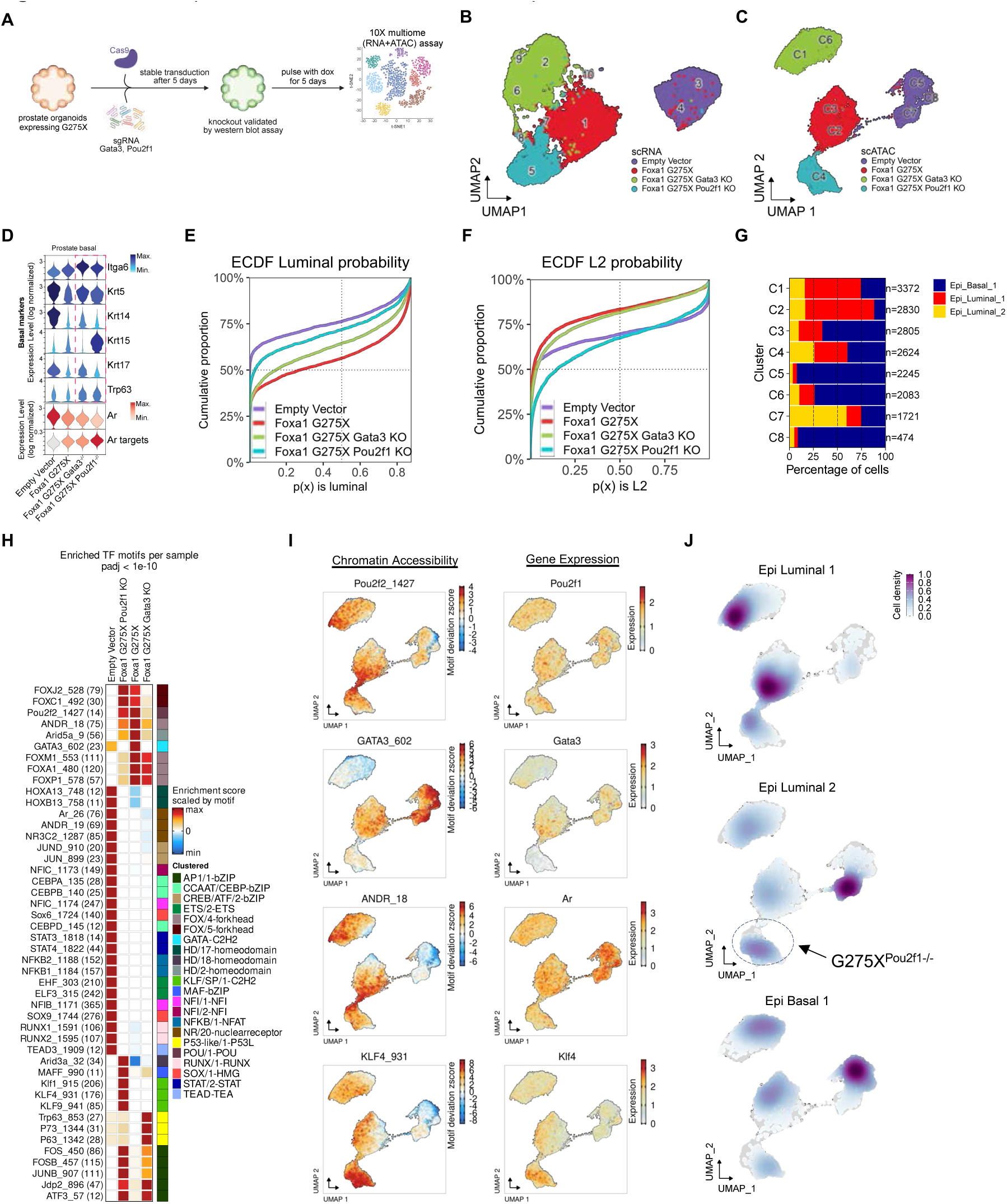
FOXA1 G275X promotes POU2F1 and GATA3 transcription factor activity to maintain luminal identity. (**A**) Experimental design schematic for organoid data generated in control and Foxa1 G275X after CRISPR knockout of Pou2f1 or Gata3 genes and used throughout this figure. After stable Cas9 and gRNA transduction, and knockout validation by western blot, organoids were pulsed for 5 days with doxycycline and subjected to 10X multiome sequencing. (**B**) scRNA-seq UMAP of the 4 genotypes: EV, G275X, G275X *Gata3-/-*, G275X *Pou2f1-/-*. (**C**) scATAC-seq UMAP of multiome samples. (**D**) Violin plots show increased expression in basal markers of G275X *Gata3-/-* and G275X *Pou2f1-/-* compared to G275X (surrounded with dotted line). *Ar* expression decreased in KOs compared to G275X, however AR targets expression increased (bottom in red). **(E)** ECDF of luminal probabilities in G275X, G275X knockouts, and EV cells. Lower lines indicate higher luminal probabilities showing most to least probable luminal cells are G275X, G275X *Gata3* KO, G275X *Pou2f1* KO, EV. (**F**) ECDF of L2 probabilities in G275X, G275X knockouts, and EV cells. Lower lines indicate higher L2 probabilities showing most to least probable luminal cells are G275X *Pou2f1* KO, EV, G275X *Gata3* KO, G275X. (**G**) Barplot of celltype abundance per cluster. (**H**) Heatmap of motif accessibilities from scATAC-seq presented by genotype. Enrichment score is based on chromvar accessibility. (**I**) UMAPs of chromVAR accessibilities and gene expression for POU2F1/2, GATA3, ANDR_18, and KLF4. (**J**) Three density UMAPS indicating highest concentration of cells for L1, L2, and Basal like cells. L2 type cells increase evident in G275X *Pou2f1* KO cells.

As expected from our initial analysis of all mutants (**Fig 1A, 2A**), UMAP projections of the scRNA-seq as well as the scATAC-seq data revealed clear separation between EV and G275X (**Fig 4B-C).** CRISPR deletion of either *Pou2f1* or *Gata3* resulted in substantial shifts of the parental G275X RNA-seq and ATAC-seq clusters, implicating both TFs in shaping the G275X-driven transcriptome. Curiously, *Gata3* mRNA levels were nearly absent in G275X *Pou2f1* KO cells, suggesting POU2F1 regulates *Gata3* expression and consistent with the presence of two POU2F1 binding peaks at the *Gata3* locus in our ChIP-seq data (**Fig S4H**). To explore the gene programs responsible for these UMAP shifts, we noted increased expression of basal-associated genes (*Itga6*, *Krt5*, *Trp63*) following *Pou2f1* or *Gata3* KO, as well as striking upregulation of *Krt15* following *Pou2f1* deletion (**Fig 4D**). Signature analysis using ECDF plots confirmed the prior finding that G275X increases the probability of L1 fate specification (**Fig 4E-F)**. This signal was mitigated by deletion of either *Gata3* or *Pou2f1*. Moreover, in the case of *Pou2f1* deletion, the reduction in L1 fate specification was replaced by a shift toward the L2 fate (**Fig 4D-F**). This shift from L1 toward L2 fate following *Pou2f1* loss was also seen in the chromatin accessibility data (**Fig S4I**).

We next examined the consequences of *Pou2f1* and *Gata3* deletion on the motifs specifically enriched in G275X organoids. *Pou2f1-/-* cells lost accessibility of the FOXA1 motif implicating POU2F1 as a key TF responsible for sustaining the Foxa1 cistrome crucial in driving the transcriptome of L1 cells (**Fig 4H-I, Fig S4J**). We also observed a marked increase in motif accessibility for KLF4, a Kruppel family and Yamanaka factor, in G275X cells following *Pou2f1* deletion, a noteworthy finding because KLF motifs are selectively enriched in L2 cells (**Fig. 4I**). GATA3 accessibility was lost in *Gata3* KO cells, as expected, but also in *Pou2f1* KO cells likely due to loss of *Gata3* mRNA expression. However, the changes in *Foxa1* and ANDR_18 seen following *Pou2f1* deletion were not seen in *Gata3-/-* cells (**Fig 4H**). *Gata3* deletion also resulted in gain of accessibility for the basal TRP63 motif, consistent with the established role of GATA3 in luminal fate specification. Taken together, our results implicate POU2F1 as a key effector of L1 luminal identity, based on loss of FOXA1 and ANDR_18 motifs and gain of L2-associated KLF family TF motifs upon *Pou2f1* deletion. POU2F1 also appears to operate upstream of *Gata3*, but the role of GATA3 in lineage specification appears to be pan-luminal, as evidenced by increased accessibility of basal TF motifs (TRP63) and loss of L1 and L2 transcriptomes following *Gata3* deletion.

### G275X has a shorter chromatin residence time and induces luminal identity faster than WT

A previous study of mutant FOXA1 alleles found that Wing 2 mutants have shorter chromatin residence times compared to WT based on fluorescence recovery after photobleaching (FRAP) and single-particle tracking (SPT) assays (*2*). One caveat of this earlier work was the use of overexpressed FOXA1 alleles, which can lead to non-physiologic effects, particularly with TFs. To determine if the G275X mutant also has a shorter chromatin dwell time, we used Cas9-mediated genome editing to insert HaloTag into the endogenous FOXA1 locus of LNCaP-AR prostate cancer cells and then derived several clones expressing either FOXA1 WT-Halo or G275X-Halo (**Fig S5C**). SPT trajectories were extracted and subject to a nonparametric Bayesian inference routine (*25*) to estimate the proportion of molecules bound to chromatin versus freely diffusing throughout the nucleoplasm. We found that substantially less G275X is associated to chromatin than WT. Similarly, G275X recovered significantly quicker than WT in FRAP experiments (**Fig S5A-B**). Consistent with earlier work on other Wing2 mutants, these results show that G275X has a shorter dwell time than WT (a so called “fast” mutant) (*2*).

To explore whether the “fast’ phenotype might be linked to the L1 lineage specification phenotype, we performed a time course experiment to ask whether the L1 signature is activated more quickly following G275X versus WT Foxa1 induction by comparing scMultiome profiles at 24 hours versus 5 days. Luminal cell probabilities determined by ECDF of scRNA-seq data remained stable in EV organoids throughout the time course but progressively increased from day 1 to day 5 in WT organoids, as expected. In contrast, maximal luminal probability (∼75% L1+L2) was reached after just 24 hours in G275X organoids (**Fig 5A-C**). The change in scATAC-seq profile of G275X organoids visualized by UMAP was similarly fast, reaching its “destination” endpoint within 24 hours, whereas the scATAC-seq profiled of WT organoids evolved over 5 days (**Fig 5A, S5G**). Commensurate with our earlier TF motif analysis, the composite FOXA1:AR (ANDR_18) and POU2F1 motifs linked to L1 fate specification were accessible within 24 hours in G275X organoids but to a lesser extent in WT (**Fig S5H**). Thus, shorter chromatin residence time of G275X relative to WT, measured by SPT and FRAP, correlates with more rapid chromatin accessibility and transcriptomic changes. While this may seem counterintuitive, there is growing evidence that TF residence time measured by SPT is often short (seconds) followed by a transcriptional burst seconds later whereas the sequence of downstream transcriptional changes required for lineage specification takes much longer (*26*, *27*).

**Fig. 5.**
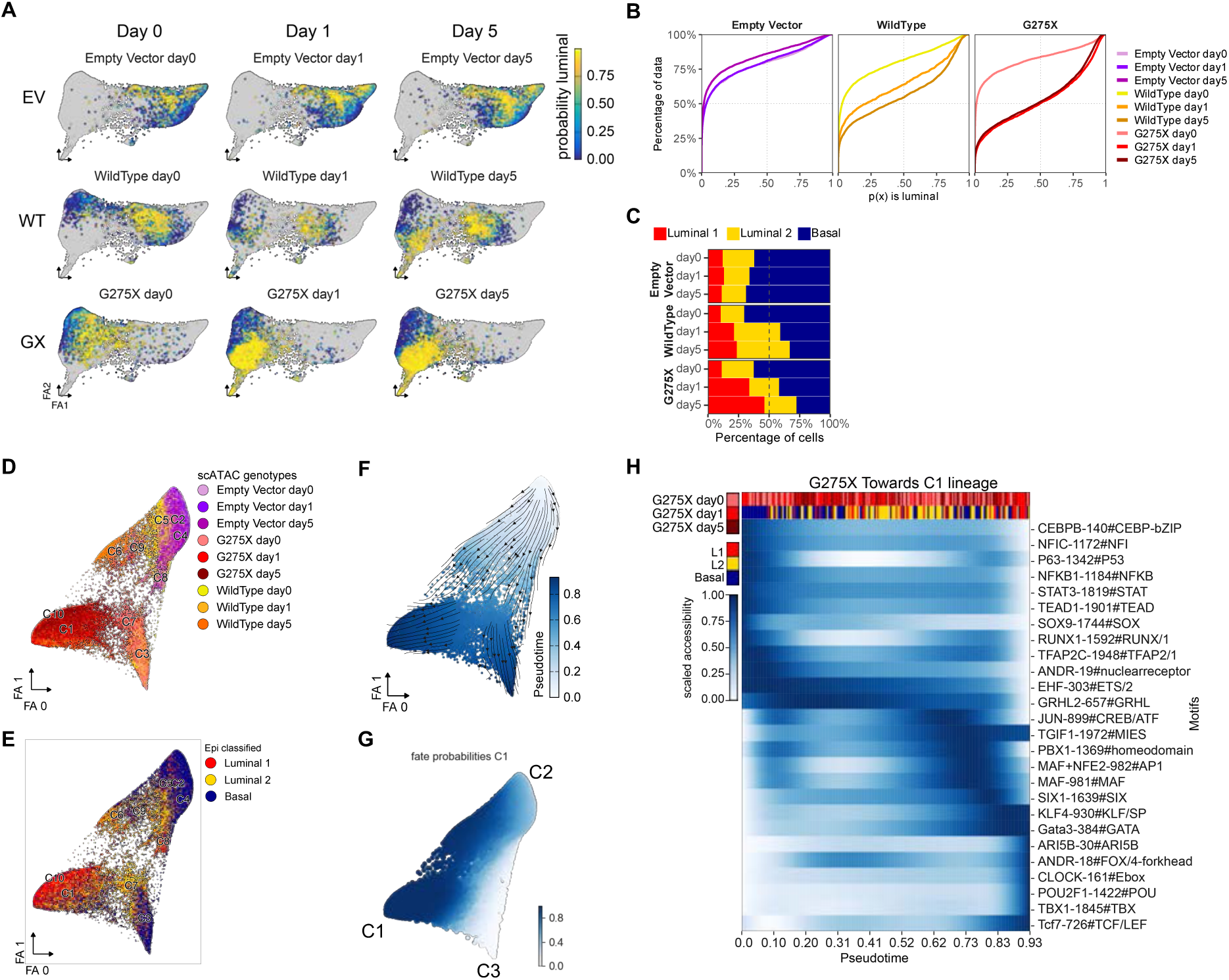
G275X mutant leads to faster activation of L1 luminal identity. (**A**) UMAPs shown by day and genotype with each cell colored by luminal probability. (**B**) ECDF curves of day0, day1, and day5 grouped by genotype. Empty Vector curves show consistent and lower probability of luminal like cells, while WT and G275X day1 curves show increased luminal cell probability. (**C**) Barplot of epithelial cell type by day and genotype showing L1 increase in WT and G275X starting at day 1 with greater increase in G275X. (**D**) scATAC-seq UMAP of cells colored by genotype. (**E**) scATAC-seq UMAP of cells colored as basal, L2, L1. (**F**) scATAC-seq UMAP colored by pseudotime using Palantir. Top corner is near the start and bottom left and right corners are two end points. (**G**) Fate probabilities as a function of pseudotime toward two branches (C3 and C1). (**H**) Heatmap of smoothed motif accessibilities within cells ordered by increasing pseudotime estimates. The first row of the Heatmap shows G275X cells colored by timepoint (day0, day1, day5) in light red, red, and dark red, respectively. The second row is colored by epithelial cell annotation with basal (blue), L2 (yellow), L1 (red). Motif names are along the y-axis with darker blue color denoting increased motif accessibility.

To investigate the sequence of TF activities required to establish luminal identity, we performed pseudotime analysis across the organoid time series (**Fig 5F**). This revealed a predominance of basal cells at day 0 (cluster C2, defined primarily by EV cells), followed by bifurcation into two branches representing luminal (L1) cells (cluster C1, primarily G275X and WT cells at days 1 and 5) and a basal-like cluster (C3 seen in WT and G275X) (**Fig 5G**). Next, we calculated and plotted smoothed pseudotime estimates for the TF motifs enriched in L1 cell fate (ANDR_18, POU2F1, GATA3 and ARID5B) (**Fig 5H**). The results show that ANDR_18 and GATA3 motifs appear early in the L1 fate transition, coincident with loss of TRP63 motifs. POU2F1 appears somewhat later, suggesting it contributes to the specification of L1 cells from L2 cells. Later in pseudotime, GATA3 motif accessibility increases further, consistent with our transcriptomic data suggesting that Pou2f1 functions upstream of GATA3 (Pou2f1 knockout results in loss of *Gata3* expression). Taken together, pseudotime analysis in concert with time series multiome profiling predicts sequential steps in TF activation that orchestrate the transition toward the L1 fate.

### Pro-L1 FOXA1 mutants drive luminal identity *in vivo*

The capability of FOXA1 mutants to drive secretory luminal cell lineage programs that are typically associated with terminal differentiation raises intriguing questions about their oncogenic potential, given that epithelial cancers often exhibit stem-like characteristics and impaired differentiation. To explore this unusual association, we examined the three FOXA1 alleles that induce an L1 luminal phenotype (WT, H247Y, and G275X) in tumorigenicity assays.

Our previous studies demonstrated that several FOXA1 mutations promote tumorigenesis in a subcutaneous model, but only in the context of *Pten* loss (*1*). Here we expanded this earlier observation but now using prostate organoids with combined *Pten*/*Trp53* deletion, a background that generates basal-like (Ck5+) tumors upon orthotopic transplantation, thereby allowing us to evaluate the luminal-inducing effects of FOXA1 mutants *in vivo* (**Fig 6A**).

**Fig 6.**
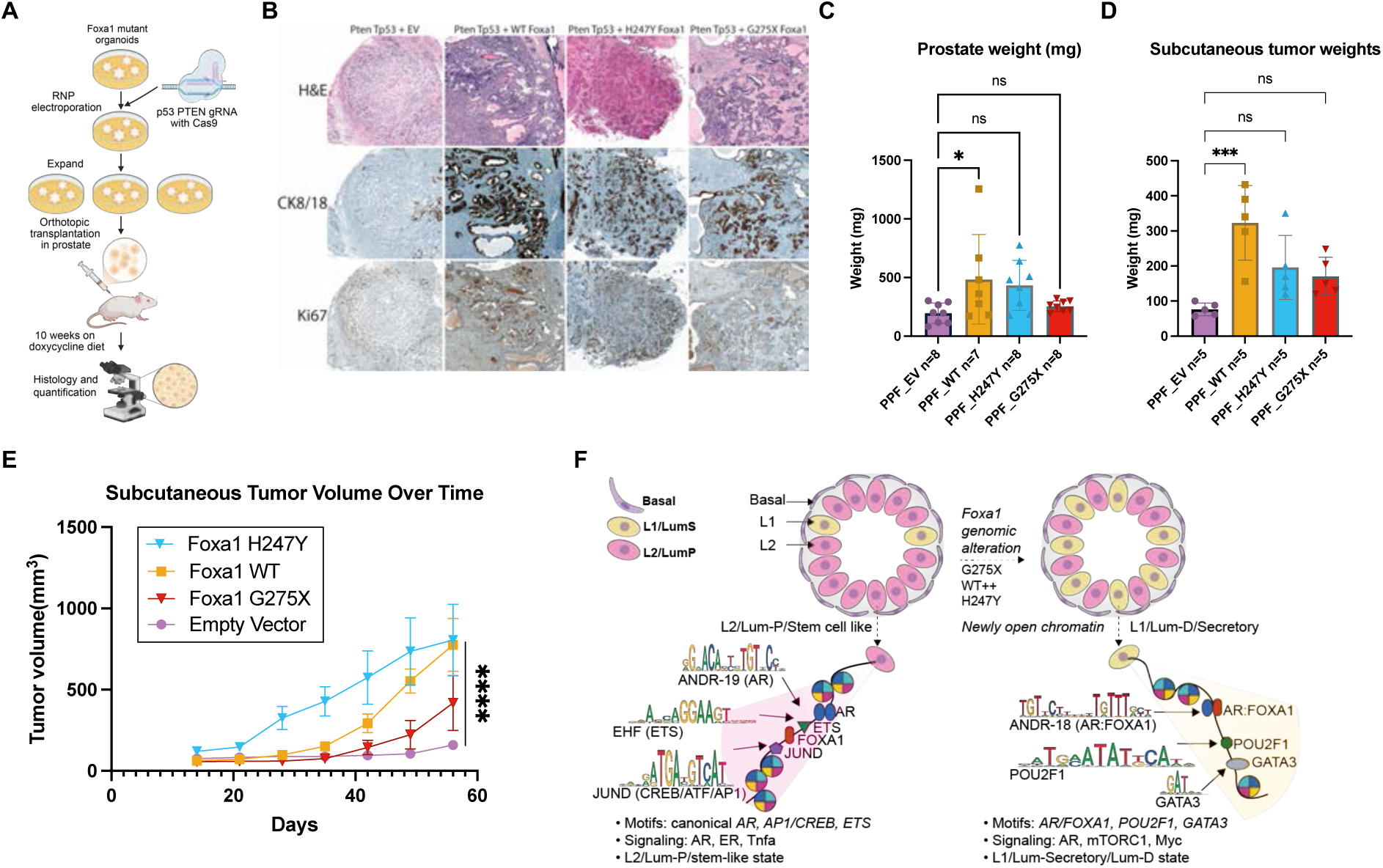
Wing2 mutants boost tumorigenicity while maintaining luminal identity. (**A**) Experimental schematic of Foxa1 mutant, *Pten*/*Trp53* KO organoids generation. These organoids were used for in vivo transplantation, subcutaneous in the right flank and orthotopic inside the prostate. The mice were maintained on doxycycline diet for the 10-week duration of the in vivo experiment. (**B**) Histology from orthotopic tumors of mice injected with mutant Foxa1-expressing organoids (WT, H247Y, G275X) display robust Ck8/18+ expression, mirroring the luminal adenocarcinoma phenotype seen in FOXA1-mutant human cancers. (**C**) Prostate weight comparisons of mutant genotypes: WT, H247Y, G275X compared to EV. (**D**) Subcutaneous tumor weight comparisons of mutant genotypes: WT, H247Y, G275X compared to EV. (**E**) Tumor volume (mm3) as a function of time in days for each of the following four genotypes: H247Y, WT, G275X, EV. Significance of interaction term (P<0.0001) was assessed by 2-way ANOVA. (**F**) Hypothetical model showing prostate organoid cell types within control and FOXA1 mutants. FOXA1 mutant organoids harboring WT overexpression, H247Y, and G275X (right) contain increasing number of L1 over L2 cells when compared to control (left). L1 cells enrich for open chromatin containing an AR:FOXA1 composite motif, POU2F1 and GATA3 motifs. Both AR signaling and mTORC1 gene expression are upregulated. L2 cells display chromatin accessible regions containing canonical AR, AP1 and ETS TF motifs as well as expression of AR targets and TNFa.

In mice injected with EV organoids (*Pten*^−/−^; *Trp53*^−/−^), invasive tumors with poorly differentiated, basal-like (Ck5+; Ck8/18−) features developed within 10 weeks. In contrast, organoids expressing either WT, H247Y, or G275X FoxA1 produced tumors with moderate-to-well differentiated histology and strong Ck8/18+ expression, closely resembling luminal adenocarcinomas observed in FOXA1-mutant human cancers (**Fig 6B**). The tumors with WT or H247Y FOXA1 also had evidence of increased proliferation (Ki67) and resulted in greater prostate weights (**Fig 6C**). Similar findings were observed in subcutaneous implantation models, where all three FOXA1 alleles accelerated tumor growth while driving luminal differentiation (**Fig 6D-E**). In summary, FOXA1 mutations that specify L1 luminal differentiation in vitro display similar activity *in vivo*, converting the basal-like histology seen with *Pten*/*Trp53* loss alone to a canonical luminal histology that resembles that seen in FOXA1-mutant human prostate cancer.

## Discussion

The high prevalence of FOXA1-mutant prostate cancer, particularly in Asian men, underscores the importance of gaining a deeper understanding of the pathobiology of these tumors and potential clinical distinctions across mutation subgroups. Through analysis of 874 such cases (2-3 times larger than previous cohorts), we largely confirm previously reported mutation patterns that now, with higher numbers, clearly segregate into 3 main subgroups: missense (Wing 2), indel/inframe (also Wing2), nonsense (truncation). A fourth well documented subgroup includes tumors with elevated levels of WT FOXA1 due to gene amplification, promoter mutations and/or structural alterations/translocations that are also common but are more challenging to identify through panel-based sequencing. Two insights not apparent in earlier cohorts include the equal frequency of nonsense (truncation) mutations in primary tumors and metastases and the fact that indels are concentrated near two residues in Wing 2 (M253, F254; 31% of all indels). Finally, access to linked clinical data from this cohort allowed us to take a preliminary look at clinical outcome that suggests nonsense (truncation) mutations may be associated with shorter survival, although this will require independent confirmation.

To explore the pathobiology across these 4 subclasses, we employed integrative single-cell analysis to unravel how mutant FOXA1 alleles drive oncogenesis. Using freshly derived mouse prostate epithelial cells, we demonstrated that mutant FOXA1 reshapes the chromatin landscape via sequence-specific binding at canonical motifs. With missense mutations (H247Y), truncations (G275X) and elevated WT (modeling FOXA1 gene amplification), AR is redirected to previously inaccessible FOXA1-AR hybrid motifs (ANDR18) and activates novel transcriptional programs that promote L1 luminal differentiation and seen in human prostate cancers with analogous FOXA1 subgroup mutations. Specifically, G275X co-opts POU2F1/OCT1 as a master regulator, establishing aberrant transcriptional circuits within days of FOXA1 expression (**Fig 6F**). Conversely, the indel mutations (M253 and FE255) activated basal-like transcriptional programs although, in this case, we were unable to confirm similar basal skewing in indel mutant human tumors (albeit with small sample size, n=17). Of note, prostates in mice engineered to express a different indel mutation (R265-71) have L1-like features (*5*), indicative of heterogeneity within subclasses depending on the specific mutation. Another variable in both studies is the reliance on mouse models to explore the transcriptomic consequences of mutant FOXA1 expression. In our case, this decision was intentional to model disease initiation in primary prostate epithelial cells rather than in prostate cancer cell lines that contain other oncogenic drivers. As technologies for culturing primary human prostate epithelial cells improve, it will be important to perform similar experiments in human models.

The discovery of the FOXA1-AR hybrid motif (ANDR18) provides a potential mechanism for how FOXA1 might reprogram AR activity and helps reconcile earlier conflicting reports of FOXA1 as both an activator and inhibitor of AR signaling (*28*), (*29*), (*30*). These discrepancies likely stem from AR cistrome repositioning from the canonical ANDR19 motif to hybrid motifs like ANDR18. ANDR18-enriched AR enhancers were also a prominent feature in the pro-L1 tumors that developed in the FOXA1 R265-71 mouse model mentioned above (*5*) as well as in a recent study of NSD2 in AR/FOXA1-dependent enhancers (*31*). Whether the changes in lineage specification associated with the FOXA1/AR hybrid motif is driven by changes in AR activity requires further functional investigation.

Our integration of single-cell multiomics and ChIP-seq with a time course experiment to track the evolution of lineage state change, coupled with computational tools such as SCARlink, enabled us to rapidly and accurately identify cooperating TF circuitry, then validate through functional studies. The identification of POU2F1/OCT1 as a key effector of mutant FOXA1-driven L1 luminal lineage specification serves as an instructive example. Building on earlier work implicating POU2F1/OCT1 and other TFs in AR signaling (*20*), the single cell approach now reveals the precise TF circuitry underlying L1 lineage fate specification. Mutant FOXA1 establishes an aberrant AR/FOXA1 cistrome (through binding to a hybrid FOXA1-AR motif), with subsequent POU2F1/OCT1 activation. It will be of interest in subsequent studies to investigate other highly ranked FOXA1-cooperative TFs, such as CLOCK, ARID5B, and KLF4, through approaches like Perturb-seq.

Whereas misssense and indel mutations all map to the Wing2 domain, nonsense/truncation mutations span the entire C-terminus downstream of the Forkhead domain, suggesting there may be greater heterogeneity within this subclass. Indeed, the pronounced L1 luminal phenotype seen with the truncation mutation examined in our work (G275X) contrasts with a different C-terminal truncation (P358fs) which induces a stem-like (L2) phenotype, underscoring the need for a comprehensive evaluation of different truncation mutants to better understand the pathobiology. This gains more relevance when considering the preliminary evidence of shorter survival of patients in this subgroup relative to those with missense mutations.

Interestingly, the L1 luminal differentiation program driven by many of these oncogenic FOXA1 mutations contrasts with the stem-like programs typically associated with epithelial malignancies. Luminal differentiation programs are also activated in ERG-translocated prostate cancers (*32*) and SPOP-mutant tumors, which are accompanied by elevated PSA expression and enhanced sensitivity to anti-AR pathway therapy (*33*), (*34*). Together with ERG, and SPOP, which collectively account for 70–80% of prostate cancers, a substantial fraction of FOXA1 mutants share a unifying theme of activating pro-luminal fate specification programs coupled with oncogenicity.

## Acknowledgements

We thank the Sawyers and Leslie Labs for valuable critiques and discussions. We are grateful to the Single Cell Analytics Innovation Lab Core Facility for help with single cell library preparation, the Integrated Genomics Operation Core Facility for next-generation sequencing,

## Author contributions

Conceptualization: EML, AN, CLS, CSL

Methodology: EML, AN, CSL, CSL

Investigation: EML, AN, TP, VBF

Formal analysis: EML, TP, VBF

Funding acquisition: CLS, CSL

Software: EML, TP, VBF, CSL

Supervision: RT, CSL, CLS

Writing – original draft: EML, CSL, CLS

Writing – review & editing: EML, CSL, CLS

## Data and materials availability

Raw and processed single cell data is available in GEO under accession number GSE285574: https://www.ncbi.nlm.nih.gov/geo/query/acc.cgi?acc=GSE285574

All other data are available in the main text or supplementary materials.

## List of Supplementary Materials

Materials and Methods

Figs. S1 to S5

Tables S1 to S3

## Supplementary Materials for

### Materials and Methods

#### Human cohort of FOXA1-mutant prostate cancer

Prostate cancer data containing **FOXA1** mutations were obtained from cbioportal.mskcc.org. After filtering for relevant cases and mutation types, the dataset comprised **991 mutations** (primary = 686; metastasis = 305) across **906 cases** from **874 patients**. The filtered dataset is provided in **Table S1**.

#### CPGEA Cohort

Normalized RNA-seq data from the CPGEA cohort was obtained and analyzed for mutation categories and expression of FOXA1. **Table S3** contains mutation and Gene expression scores.

#### LnCAP-AR cell culture and endogenous Foxa1-HaloTag knock-in

LnCAP-AR cells (*36*) were cultured with RPMI-1640 with 9.1% FBS with penicillin and streptomycin and kept in a water-jacketed incubator at 37°C with 5% CO2. Cells were generally split 1:8 every three days with 0.25% Trypsin to facilitate detachment.

HaloTag was knocked-in at the endogenous FOXA1 locus with plasmid-encoded SpyCas9, sgRNAs, and homology directed repair templates. For WT Foxa1, the sgRNA targeted the C-terminus of the protein. For Foxa1-truncation, the sgRNA targeted the ORF where the truncation occurs. In both cases, repair templates containing a linker and HaloTag were cloned with ∼500 bp upstream and downstream homology to the cut site **[Plasmids are available upon reasonable request.].** The following Cas9 protospacer-PAM sequences were used for endogenous HaloTag editing:

WT-guide: CGGGTCTGGaatacacacct[tgg]
GX-guide: caagtgcgagaagcAGCCGG[GGG]

LnCAP-AR cells were electroporated with all plasmids above using a homemade buffer (*37*) and the T-009 protocol of the Lonza Nucleofector II. Single clones were isolated by FACS and sequence-validated to have the by PCR and Sanger sequencing. Clones used for this study were also Western blotted for Foxa1. The cell lines tested negative for mycoplasma about monthly by PCR and were confirmed to be LnCAP cells by STR profiling.

#### Single particle tracking at fast framerates (fast SPT)

SPT experiments were conducted on a custom-built Nikon TI microscope with a 100x/NA 1.49 oil-immersion objective to allow for Highly-inclined Laminated optical (HiLo) illumination (*38*).

About 18-24 hours prior to imaging, LnCAP-AR cells were plated on MatTek #1.5 glass-bottom dishes by first coating the dishes in 0.01% poly-L-lysine and rinsing with distilled water. The day of imaging, cells were stained for ∼5 minutes with 1 nM JFX650-HaloTag ligand and 10 nM JFX549-HaloTag ligand (*39*), then rinsed with pre-warmed media and incubated in more media for 20-30 minutes to allow for unbound dye to wash out. The cells were then immediately imaged, during which they were kept in an incubation chamber maintaining a measured 37°C 5% CO2 environment.

Cells were located using a 561 nm laser and imaged with the 635 nm laser in the following way for 10000 cycles: 7.48 ms frame interval (7 ms and a 0.48 ms camera transition time), 1 ms pulsed 635 nm laser powered to 1100 mW at the beginning of the camera integration phase, 0.48 ms 405 nm laser during the camera transition time to reactivate dyes that have entered a dark state. 13-22 cells of each clonal line were imaged each day and repeated over 3 days; one day’s results are shown, though the data are very reproducible day-to-day.

Spots from movies were detected, subpixel-localized, and connected by quot (https://github.com/alecheckert/quot), and analyzed using saspt (https://github.com/alecheckert/saspt; (*25*), a published open-source method that infers proportions of diffusive states using Bayesian inference. For ease of comparison, we sum and highlight the inferred proportion of molecules diffusing below 0.1 μm2 s-1 **[Trajectories are available upon reasonable request.]**

#### Fluorescence recovery after photobleaching (FRAP)

FRAP was performed on clonal LnCAP-AR lines, as well as the parental LnCAP-AR line exogenously expressing H2B-HaloTag as a control. Cells were stained with 50 nM TMR-HaloTag ligand for 5 minutes, then rinsed with pre-warmed media and incubated in more media for 20-30 minutes to allow for unbound dye to wash out. The cells were then immediately imaged, during which they were kept in an incubation chamber maintaining a measured 37°C 5% CO2 environment.

Cells were imaged on a ZEISS LSM 900 in confocal mode using a 63x oil-immersion objective using the 561 nm laser for acquisition. A 2 μm diameter circle was bleached with both the 561 nm and 405 nm lasers after 20 frames of initial acquisition, and the cell was imaged for ∼150 seconds afterwards to track recovery. The bleach spot was drift-corrected after imaging; recovery of the spot was divided by the intensity of a frame-matched nuclear mask.

#### Organoid culture

All cell lines and organoids were periodically tested and found negative for mycoplasma test (Lonza #LT07-318). Cell lines used in this study were maintained in a 37 ^ο^C and 5% CO2 incubator. Murine organoids were established from adult mouse prostate taken from C57/BL6 mouse, as previously described (*11*). Briefly, entire mouse prostate was harvested, and physically dissociated into smaller pieces, with sterile razor blades. Next, it was digested with collagenase type II (Gibco) for about 1hr at 37 ^ο^C with gentle shaking. Next, TrypLE (Gibco) was added for digestion at 37 ^ο^C until a single-cell suspension was obtained. Single cell suspension was washed 3 times with cold PBS after TrypLE digestion. Organoid media was prepared as previously described (*11*). It was freshly supplemented with Y-27632 (10uM) to inhibit anoikis. Finally, cells were filtered through FACS tubes and seeded for organoid culture. Bulk isolated prostatic epithelial cells were embedded in 35-μl drops of basement membrane extract (Matrigel, Corning) and overlaid with mouse prostate organoid medium.

For transduction, organoids were transduced with lentivirus for pCW-EV and pCW-Foxa1 (WT and mutant cDNA) and were selected with 2ug/ml of puromycin for 5 days. The pCW plasmids were cloned as previously described using Flag-tagged alleles (*1*). Antibiotic selection was started 3-4 days after viral transduction. Organoid sorting and 3D culturing was performed as described previously (*1*). Transduction with Lenti-Cas9-Blast was followed by 5-days of selection in 10ug/ml of blasticidin. For most in-vitro assays assessing the effect of the Foxa1 transgene, unless otherwise mentioned, organoids were seeded and treated with doxycycline at 500ng/ml to induce the expression of the Foxa1-2A-dsRED fusion cDNA and were harvested 5 days after the addition of doxycycline.

#### Lentivirus production and transduction

Human HEK293 cells (ATCC. CRL-1573) were grown in DMEM (ATCC. 30-2003) with 10% FBS, 100 U/mL penicillin-streptomycin and 2mM L-glutamine. Lentivirus packaging was performed in HEK293T using Lipofectamine. 2000 reagent (Invitrogen) in accordance with the manufacturer’s instructions. Medium containing virus was concentrated using PEG-it Virus Precipitation Solution (System Biosciences). Lentiviral transduction was performed as described previously (*11*).

#### *Pten*/*Trp53* knock out in organoids

*Pten* and *Trp53* knock outs were generated by electroporation of Cas9-gRNA (RNP) complexes with the Lonza kit (Lonza, VVCA-1001), following instructions provided in the Lonza manual. Briefly, 1 million dissociated organoid cells were resuspended with nucleofection buffer, RNP complexes, and electroporation enhancer (IDT 1075915, 1:1molar ratio to cRNP) in a total volume of 100 μL. The cell suspension was transferred to a nucleofection cuvette and nucleofected using Lonza Amaxa Nucleofector II (program T-030). Cells were centrifuged at 400g for 5min and seeded for culture in matrigel, with organoid media. All guide RNAs used for the study were generated using the CRISPR Design Tool (http://crispr.mit.edu). Following gRNA sequences were used for the study:

*Pten* guide: ACCGCCAAATTTAACTGCAG
*Trp53* guide: ACCCTGTCACCGAGACCCC
non-targeting guide: CTTCACGCCTTGGACCGATA

#### *Gata3* and *Pou2f1* KO in organoids

*Gata3* and *Pou2f1* gRNAs were cloned into lenti-lvt-pUSEG-GFP plasmid. Cas9 was transduced with lentiCas9-blast (addgene plasmid 52962). The organoids were infected with Cas9-blast and gRNA virus and selected with 2ug/ml of puromycin and 10ug/ml of blasticidin. After 5 days of selection, KO was confirmed by western blot assay on whole cell fractions. Guide 2 for Gata3 and guide 1 for *Pou2f1* were more efficient in knocking out their respective genes. These lines were subsequently used in multiomic assays. All guide RNAs used for the study were generated using the CRISPR Design Tool (http://crispr.mit.edu). Following gRNA sequences were used for the study:

*Gata3* guide 1: GGGACACGATCCTCAGCACA
*Gata3* guide 2: GTTGCAGTTTCCTTGTGCTG
*Pou2f1* guide 1: TCCCGTTCCTTCCTCTCCCG
*Pou2f1* guide 2: TCGGGAGAGGAAGGAACGGG
non-targeting guide: CTTCACGCCTTGGACCGATA

#### In vivo mouse experiments

All animals (mice) used in the study were cared for in accordance with guidelines approved by the Memorial Sloan Kettering Cancer Center (MSKCC) Institutional Animal Care and Use Committee (IACUC) and Research Animal Resource Center (RARC). For orthotopic transplants, NOD SCID Gamma (NSG) mice were purchased from Jackson laboratories, and 1 million cells were injected into the dorsal lobe of the prostate in 8-week-old mice. Cells were prepared after dissociating 3D organoids into single cell suspension. For each mouse, the appropriate number of cells were resuspended in 10ul of matrigel and 10ul of organoid media, a total volume of 20ul. For sub-cutaneous injections, organoids were dissociated, and 1 million cells were resuspended in matrigel and media, in 1:1 ratio. 20ul of cell suspension was then implanted in the right flank of a NOD SCID Gamma (NSG) mice were purchased from Jackson laboratories.

#### Immunohistochemistry (IHC) and immunofluorescence (IF)

H and E staining and IHC was carried out by the MSKCC Molecular Cytology Core using Ventana Roche Benchmark Ultra and Leica. For tissue fixation, murine tissue was fixed using 4% paraformaldehyde overnight, dehydrated with ethanol, and paraffin embedded per standard protocol. 4-μm slides were cut and placed on glass slides. Immunohistochemistry or immunofluorescence of FFPE tissue was performed using a Ventana BenchMark Ultra and Leica. FFPE stained tissue was scanned using a MIRAX scanner and processed using caseviewer software. The antibodies used are CK8/18 abcam, ab53280 (1 to 1000 dilution), and Ki67 cst, 12202T (1 to 500 dilution).

For IF staining of wholemount organoids, organoids were fixed with 4% PFA on ice for 1 h, then blocked and permeabilized using 1% triton-X100 and 1% FCS in PBS0 for 1 hour at RT. Staining was performed in 0.3% Triton X-100 and 0.0.5% FCS in PBS overnight at 4 degrees on gently shaking. Stained organoids were imaged using a Leica SP5 or SP8 confocal microscope. Images were processed using Leica software or FIJI software.

#### Western blot assay and analysis

Organoids were isolated from the matrigel basement membrane extract by typle E (trypsin) treatment and performing multiple washes with ice cold PBS in a 15-ml Falcon tube. Cells were lysed in RIPA buffer containing protease inhibitors (Calbiochem) and phosphatase inhibitors (Calbiochem) on ice and sonicated three times for 30 s at 30-s intervals using a Bioruptor. Lysate were centrifuged for 10 min at 20,000G at 4deg. The supernatant was collected as protein lysate. Protein concentrations were quantified using a bicinchoninic acid (BCA) assay (Pierce, Thermo Fisher). Lysates were denatured using 4X protein loading dye (SDS 200 nM Tris, 8% SDS, 0.4% bromophenol blue, 40% glycerol, 400 mM 2-mercaptoethanol, pH 6.8). 10-20 μg of protein was loaded on NuPage 4-12% gradient bis-tris polyacrylamide gels (Invitrogen). After electrophoresis, protein was transferred to a PVDF membrane and blocked with either 5% milk or 5% BSA in TBS-T. Primary antibodies were incubated overnight. Membranes were washed using TBS-T and incubated with secondary antibodies for 1 h at room temperature. Proteins were visualized using ECL and ECL prime (Amersham, GE healthcare) and ImageQuant 800 (Amersham, GE healthcare). Blots were analyzed using FIJI/ImageJ software. The following antibodies were used in the assay, FOXA1 sigma, SAB2100835-100UL (1 to 1000 dilution), Vinculin cst, 13901S (1 to 5000 dilution), Pou2f1 abcam, ab15112 (1 to 1000 dilution), Gata3 abcam, ab199428 (1 to 1000 dilution), Cyclophillin B cst, 43603S (1 to 5000 dilution), and TBP abcam, ab63766 (1 to 5000 dilution).

### Processing of single cell RNA sequencing

#### Read alignment of single nucleus RNA sequencing

Single-cell RNA sequencing data (scRNA-seq) was first processed using 10x Cell Ranger ARC software (ref), which includes alignment and minimal cell filtering for both scRNA-seq and scATAC-seq. This pipeline was customized to align sequencing reads to the Mus musculus genome (mm10) plus an additional sequence for the Foxa1 DsRed reporter.

#### Filtering expression matrix

The ARC filtered count matrix was examined with output plots to visualize cell and gene coverage distributions, and mitochondrial and ribosomal counts (**Fig. S1L**). Based on these plots several quality metrics were used to remove poor quality cells (those with fewer than 500 transcripts) or putative multiplets (defined as cells containing greater than 100K transcripts). Cells with greater than 40% mitochondrial reads were filtered out from further consideration.

#### Expression matrix normalization and variance stabilization

To account for technical variation and normalize expression data the R package Seurat (*40*) was used on the filtered integer gene count matrix. The sctransform method (*41*) was used to perform normalization, variance stabilization, and feature selection. We regressed out cell cycle scores and mitochondrial percentage using vars.to.regress parameter of SCTransform function.

#### Dimensional reduction and visualization

A nearest neighbor graph of cells was learned and used as input to the Leiden algorithm for cell clustering. Dimensionality reduction and uniform manifold and approximation projection (UMAP) and force directed layout using force atlas (FA) were applied to visualize cell clusters. A total of ten different cell clustering resolutions were evaluated and scored by silhouette score, a clustering metric. UMAPs were generated and used to hypothesize cell similarities and identify clusters for differential expression testing. Quality control, filtering, normalization and clustering of all cells were performed resulting in 20825 cells. Visualization of single nuclei on a UMAP from scRNA organoids depicted genotypically distinct cell transcriptomes.

### Differential genes and pathways

Differential expression testing was performed using the R package Seurat with presto (*42*). Genes were ranked using the average log foldchange and a false discovery rate adjusted p-value. Differential expression of L1, L2, and Basal signatures compared across 6 conditions ranked by AUC were plotted (**Fig. 1H**) and can be found in **Table S2**. A GSEA analysis with fsGSEA (*43*) utilized a list of ranked genes per comparison and the collection of molecular signatures database.

### Single nucleus Assay for Transposase-Accessible Chromatin with high-throughput sequencing data (scATAC-seq)

#### Filtering features matrix

Single-cell Assay for Transposase-Accessible Chromatin with high-throughput sequencing data (scATAC-seq) was analysed using the ARC filtered count matrix.

Cell filtering, feature selection, and analyses were performed using ArchR software (*44*). We generated output plots to visualize distributions of cell counts within promoters, gene bodies, and transcription start sites (TSS). Quality metrics were used to remove poor quality cells including low TSS enrichment (<4) or low number of unique nuclear fragment (<1000) or putative multiplets (defined as cells containing greater than 1M reads).

#### Peak and motif identification

Peaks with scATAC-seq data were called using MACS2 (*45*) through the ArchR interface. Differentially accessible regions weredefined as those having significant difference with log2 fold change ≥ 0.5. For motif detection we utilized 2 databases, 1) CisBP and 2) the Non-redundant TF motif matches genome-wide (Vierstra2020). Motif matching within peaks was performed using MotifMatcher with defaults. The gene identity of each position weight matrix match was recorded so that we retained only those motifs whose corresponding gene was expressed. To identify motifs statistically significant between comparisons we calculated motif chromVAR (*46*) scores for cell and performed Spearman correlation of those scores to the motif’s gene expression values. Those motifs having positive correlation with expression and significant chromVAR scores were kept for further analysis.

### Read alignment and peaks atlas construction for chromatin immunoprecipitation by sequencing (ChIP-Seq) data sets

Alignments and peak files generation were performed with custom bash scripts that conformed to the ENCODE consortium ChIP analysis pipeline. First, sequencing reads were aligned to the Mus musculus genome, mm10 using bowtie2 (--max_fragment_length 2000 –trim-qual 15). Picard’s mark duplicates was used to remove PCR and optical duplicate reads. We filtered out lower quality alignments using samtools (samtools view -F 1804 -bSq 30).

The de-duplicated, quality filtered alignment files were input to MACS2 for peak calling with a permissive p-value cutoff ≤ 0.2 to provide sufficient signal for Irreproducible Discovery Rate (IDR) analysis. ENCODE blacklisted regions were removed from the results and the output saved as narrow peak files. To standardize peak length the narrow peak entries were centered at peak summits and flanking regions were extended by 250 nucleotides. Next, we utilized Irreproducible Discovery Rate (IDR)for between replicate reproducibility of summits with a false discovery rate (FDR) ≤ 0.01. A final atlas of peaks was constructed by combining results from the above steps.

### Differential ChIP-seq peaks and motif identification

A PCA of AR ChIP peak data confirmed replicate similarity, and luminal samples (H247Y, G275X, WT) grouped to one side while EV grouped with *Ar* KO. Differential peaks were determined by DESeq2 (*47*). Briefly, the atlas of peaks were normalized using DESeq2’s median of ratios normalization using a model design: **∼0 + condition** for a one versus others comparison. K-means was used to assess the number of peak clusters for each ChIP-seq set. ChIPseeker (*48*) was then used to annotate genomic features of the peaks(**Fig. S3A, Fig. S3B**)

Motif calling was performed within peaks by the Motif Occurrence Detection Suite (MOODS) library through the motifmatcher function in R. For motif identification we searched 2 databases, 1) CIS-BP and 2) the non-redundant TF motif matches genome-wide (Vierstra2020), containing mouse and human motifs. We considered only motifs having a p-value cutoff of 5e-05, within a discovery window of 7 nt. against the background frequency of Mus musculus genome nucleotides. To discover motifs differentially enriched we performed a binomial test of the number of significant motifs in one sample’s peaks vs. other samples (**Fig. S3C, Fig. S3E**).

POU2F1 ChIP data was aligned to mm10 genome using bowtie2 (--local_preset "--very-sensitive-local --no-mixed --no-discordant --phred33") and filtered using samtools (samtools view -F 1804 -bSq 30). Duplicates were marked using picardtools. Replicate peaks were assessed using Irreproducible discovery rate (IDR) and retained if significant at IDR score ≥ 830. The atlas of peaks were normalized and compared between conditions using DESEQ2. POU2F1 peak read counts were compared in Foxa1 G275X (n=2607) vs Empty Vector (n=4371) and assessed for significance. Peaks significantly upregulated either in Empty Vector or Foxa1 G275X (adjusted p-value < 0.05 & log2FC > 0.5) were used for motif discovery performed with MOODs library. Only motifs having a p-value cutoff of 5e-05, within a discovery window of 7 nt were considered hits. Motif enrichments were calculated using binomial z-score comparing each genotype against the background set of peaks from the scATAC-seq peak atlas. Specifically, if *p* represents the probability that a peak in the background set contains an occurrence of the motif, then the binomial Z-score for a cluster of size *N* with *C* peaks containing the motif is 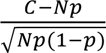. Although these Z-scores do not incorporate a correction for multiple hypotheses, in practice the top-ranked motifs have such strong enrichments they would still be highly significant after correction.

### Milo analysis

We generated pseudo-replicates per sample from our single cell expression data and performed differential abundance testing using Milo (*35*). Neighborhood creation was performed using the following parameters makeNHoods(refinement_scheme = "graph", prop = 0.1, k = k, d=30, refined = TRUE, reduced_dims = "PCA"). A design matrix to compare each mutant to the Empty Vector control was created and used as input to makeContrasts(). Neighborhood tests were performed for all model contrasts, saved and plotted to show differential neighborhoods.

### Gaussian mixture model

Single cell gene signature scores for each cell were input as a matrix to the function densityMClust() to perform point density estimation by a Gaussian finite mixture model for model assessment using the Mclust package (Scrucca L., et al.). We then used the Mclust() function to fit the data to a 2-component gaussian mixture model for luminal/basal classification. This demonstrated 2 distinct densities and allowed per cell luminal or basal probability assignment (**Fig. S1G, Fig. S1H**). We also fit a 3-component gaussian mixture model for basal, luminal1, and luminal2 probability assignment to each cell and performed dimensional reduction using MclustDR() to reduce dimensionality with a set of linear combinations, ordered by importance as quantified by associated eigenvalues of the original features, similarly to PCA. (**Fig. S1I**). A discrete cell classification was made by assignment to the class with greatest probability. We then calculated a heatmap showing the top differential genes per cell classification (basal, luminal1, luminal2) (**Fig. S1F**)

### Hotspot

To identify genes that covary in cells in an informative manner and organize these genes into modules we performed Hotspot analysis (*14*). We used the single cell knn graph as input to hotspot to compute autocorrelations and find informative genes. We then selected highly significant genes according to Hotspot having an FDR <0.01 (n=1850) and computed local correlations before grouping into modules. Gene modules were assessed using gprofiler2 and three query databases: gene ontology (GO), Kyoto Encyclopedia of genes (KEGG) and Reactome (REAC). The top pathways (within top 5) from GO were annotated on the heatmap.

### AR regulated targets

We utilized multiome data output that jointly sequences chromatin accessibility using scATAC-seq and gene expression using scRNA-seq from the same cell. For each modality cells were combined to generate an ATAC and an Expression signal corresponding to each lineage specific state (L1, L2, Basal). We then used SCARlink to predict the expression data of each gene from chromatin accessibility data by defining the genomic region around the gene body with additional 250KB flanking regions extending the gene’s transcription start and transcription end sites. These regions were segmented into 500-nt tiles and ranked using SCARlink-derived Shapley scores to indicate the importance of each tile to the prediction. For each gene the Shapley scores were converted to z-scores and adjusted for gene length and used to rank putative enhancers (tiles).

Next, we gathered Ar ChIP-seq data overlapped with significantly differential scATAC-seq peaks per lineage specific state to assign ChIP-seq peaks to a specific state (**Fig. S3F**). We then cross-referenced Ar ChIP-seq peaks, accessible AR motifs (ANDR_18 or ANDR_19) and SCARlink length adjusted z-scores to pinpoint Ar-regulated enhancers of TF genes. The overlap of these signals was converted to a score by calculating the dot product of the length adjusted z-score from SCARlink, the normalized peak counts from scATAC-seq and the ChIP normalized counts. Those loci that contained SCARlink enhancers overlapped with motifs and overlap with ChIP data were ranked by their score and plotted.

### Epithelial versus FOXA1 expression in FOXA1-mutant tumors

We analyzed processed RNA-seq data from the Chinese Prostate Genome and Epigenome Atlas (n = 134 cases with matched tumor/normal profiles and annotated FOXA1 mutation status) (*4*). Epithelial output scores were computed using gene signatures derived from normal mouse prostate (*11*). These mouse signatures were converted to human orthologs, and expression values were standardized as z-scores across all samples. Briefly, z-scores were calculated by subtracting the gene’s mean expression across all samples from each individual sample and dividing by the standard deviation of gene expression across all samples. For each sample, the sum of z-scores for L1, L2, or basal genes yielded an epithelial output score. FOXA1 expression was then plotted against each epithelial score in separate panels (**Fig. 1L**).

---

**Fig S1.**
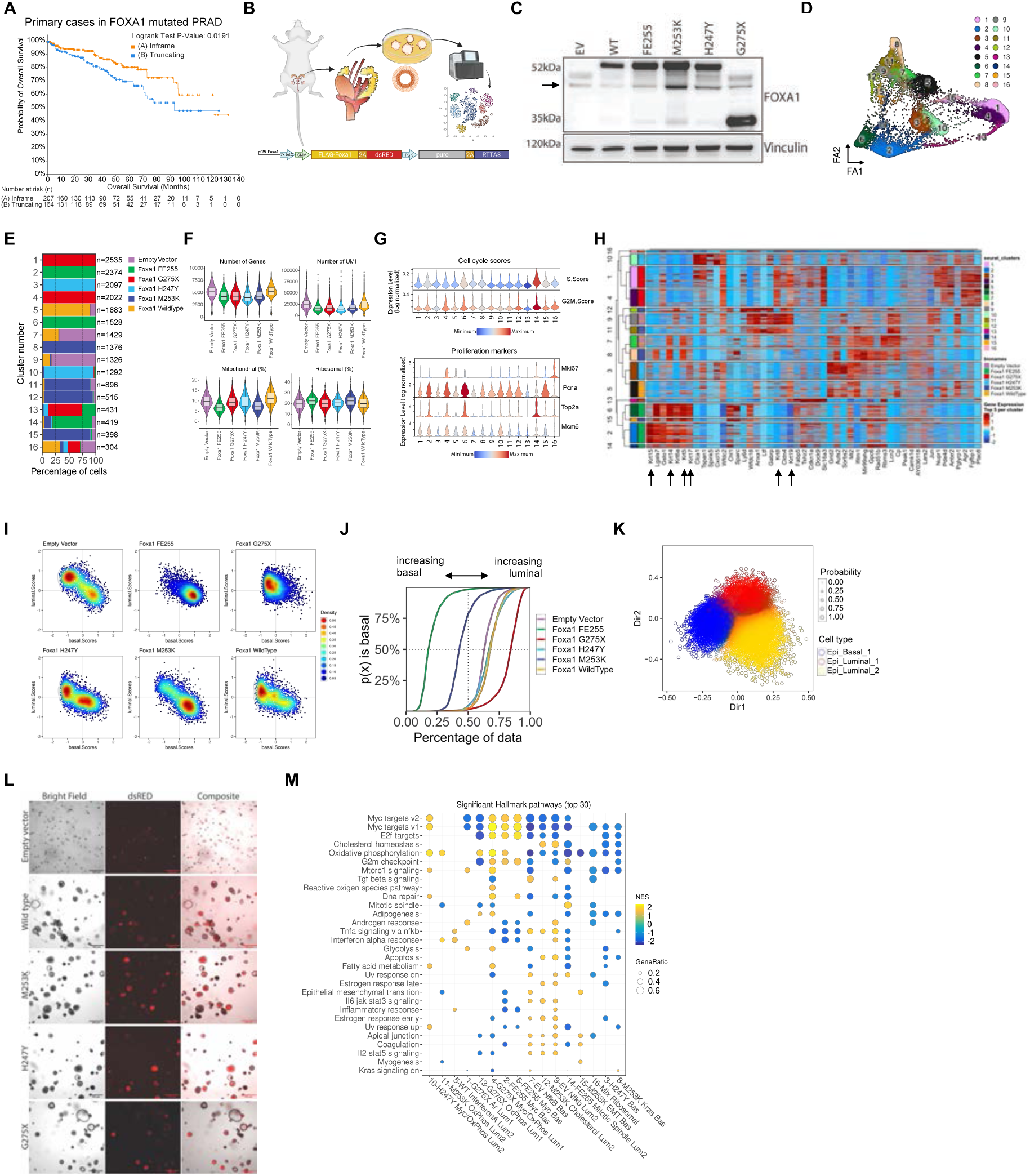
FOXA1 mutations frequently found in patient prostate tumors drive epithelial cell fate changes in prostate organoids. (**A**) Survival plot in primary prostate cases having In-Frame or Truncating FOXA1 mutation. (**B**) Experimental design of FOXA1 mutant organoids engineered from mouse prostate. (**C**) Western Blot of FOXA1 mutants used in study, all of which are Flag-tagged (see Methods). Arrow indicates endogenous FOXA1 protein. (**D**) scRNA-seq UMAP colored by Leiden clusters. (**E**) Bar plot of clusters defined by Leiden clustering of scRNA-seq data. Cluster number on left axis and number of cells annotated on right side. (**F**) Violin plots of scRNA-seq basic quality metrics used per sample. (**G**) Violin plots of Cell cycle scores (top) and proliferation markers (bottom) by cluster number. (**H**) Differential gene expression (DGE) heatmap by cell cluster showing prostate luminal (*Krt8*, *Krt19*) and basal (*Krt5*, *Krt14*, *Krt17*) cell markers among top DGE. (**I**) 2D density plots of luminal and basal scores per cell used as motivation for multivariate Gaussian fitting. (**J**) Empirical cumulative distribution function (ECDF) of basal cell probabilities (luminal prob. = 1-basal prob.) per genotype. Curves shifted to the left show higher probability of cells as basal and right shift show higher proportion of cells as luminal. (**K**) PCA-based dimensionality reduction after model-based clustering and classification. Points represent cells colored by epithelial cell type. (**L**) Organoid bright field staining of transgene in DSRed for EV, WT, M253K, H247Y, and G275X genotypes. (**M**) Hallmark pathways assessed via GSEA and shown per cell cluster.

**Fig S2.**
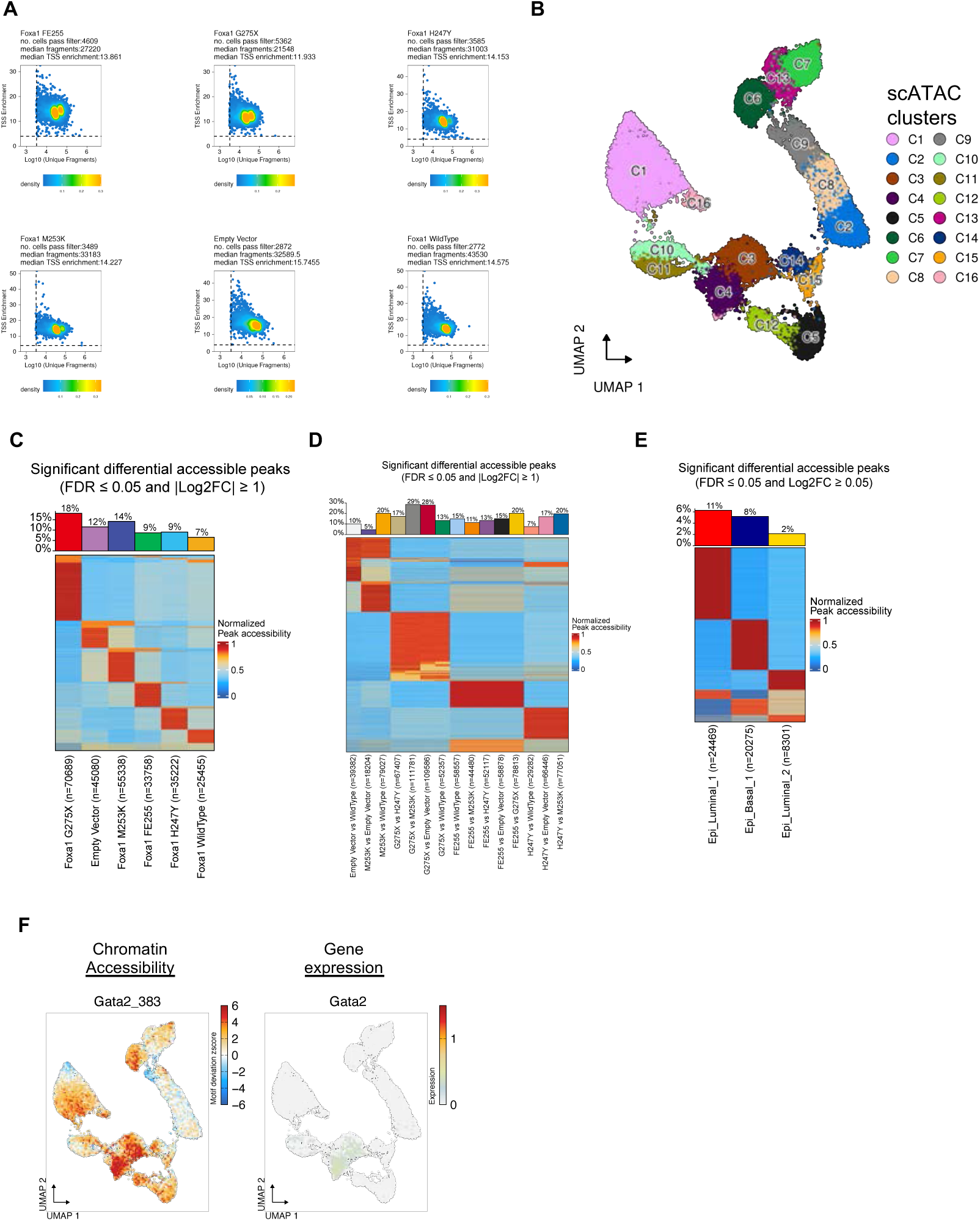
FOXA1 single nucleus chromatin accessibility reveals distinct transcription factor motifs that integrate luminal / basal expression programs. (**A**) scATAC-seq panels showing density of cells after quality filtering using Log10 (Unique fragments) on the x-axis and TSS Enrichment score (ratio of reads in TSS vs flanking regions) on the y-axis. (**B**) scATAC-seq UMAP of cells colored by leiden clustering. (**C**) Heatmap of average peak accessibilities per genotype that are significantly differential in one compared to all other genotypes (FDR ≤ 0.05 and |Log2FC| ≥ 1). (**D**) Pairwise comparison heatmap of average peak accessibilities significantly differential (FDR ≤ 0.05 and |Log2FC| ≥ 1). (**E**) Heatmap of average peak accessibilities per cell type that are significantly differential (FDR ≤ 0.05 and |Log2FC| ≥ 1). (**F**) UMAPs of GATA2 chromatin accessibility motif deviation score (left) and expression of mRNA (right).

**Fig S3.**
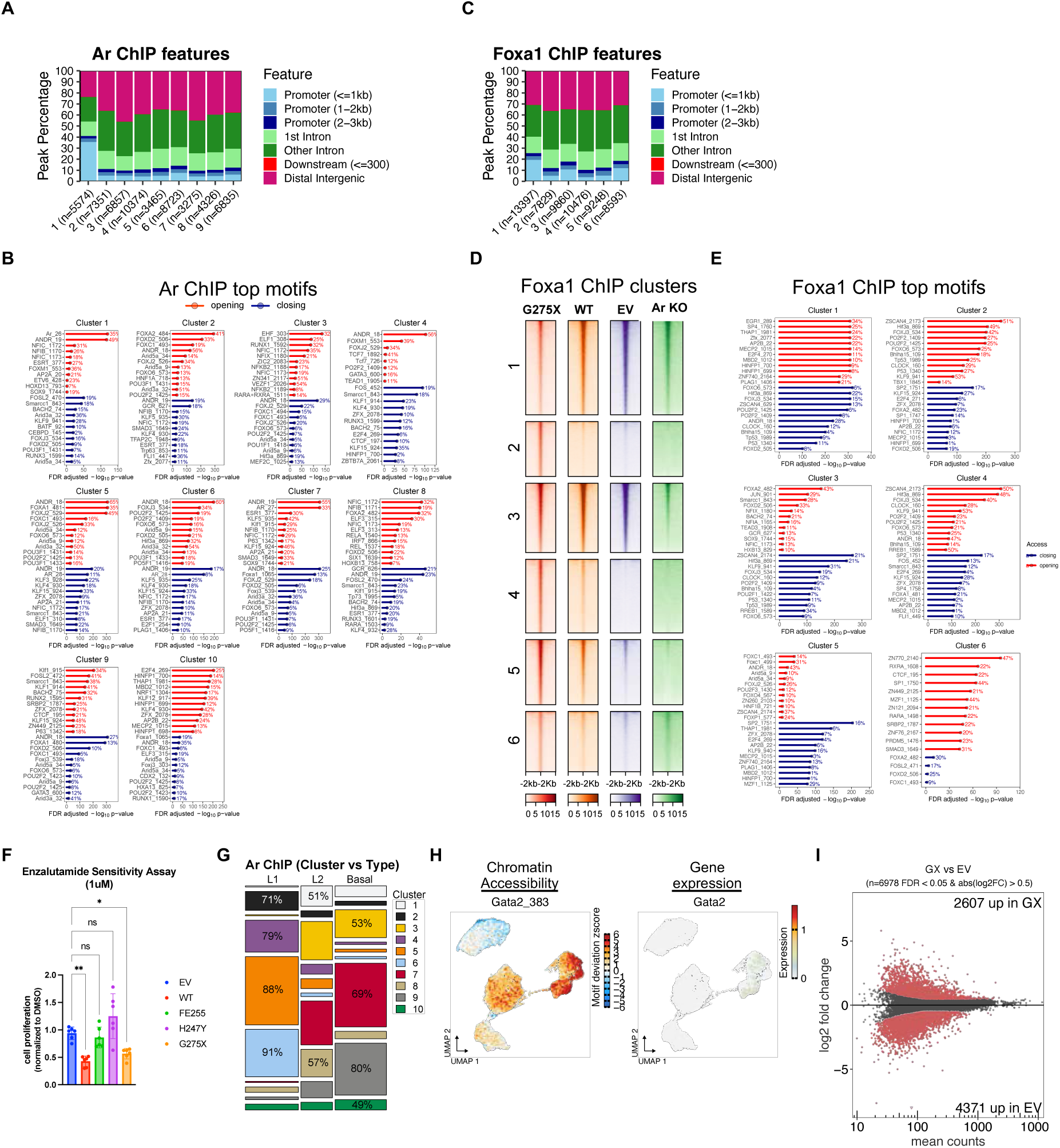
FOXA1 G275X cooperates with Androgen receptor by enriching for an alternative AR motif and reinforces AR target gene expression. (**A**) AR ChIP-seq peaks labeled by % of features. (**B**) Top motifs detected within AR ChIP-seq peaks per cluster. (**C**) FOXA1 ChIP-seq peaks labeled by % of features. (**D**) Tornado plot of FOXA1 ChIP-seq signal after K-means clustering in G275X, WT, EV, and *Ar* KO samples. (**E**) Top motifs detected within FOXA1 ChIP-seq peaks per cluster. (**F**) Organoid sensitivity measurements to 1 micromolar of the Androgen Receptor antagonist Enzalutamide. Cell proliferation measurements are shown normalized to the Empty Vector control. Pairwise statistical significance is indicated above braces. (**G**) Tile plot displaying AR ChIP-seq peaks clustered and categorized into epithelial cell types (basal, L1, L2) based on scATAC-seq overlap. Percentages representing the fraction of peaks by epithelial cell type are only sown for those significantly different between epithelial cell types. (**H**) UMAPs of GATA2 chromatin accessibility motif deviation score (left) and expression of mRNA (right). (**I**) MA plot of POU2F1 ChIP peaks. G275X significant peaks (FDR < 0.05 and |log2FC| > 0.5) are shown in red above the 0 on the yaxis. EV significant peaks (FDR < 0.05 and |log2FC| < 0.5) are shown in red with a -log2FC and below the 0 line on the y-axis.

**Fig S4.**
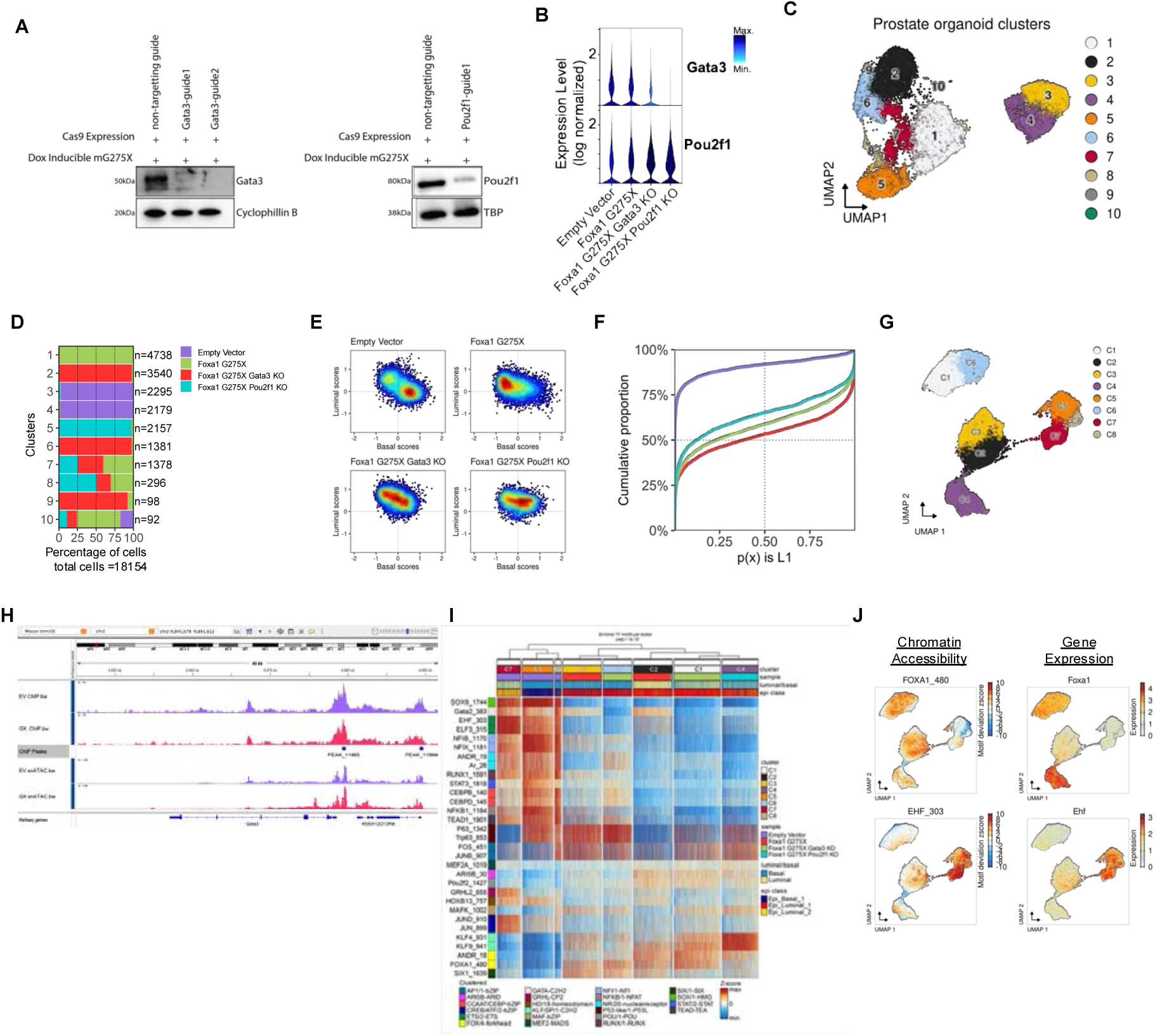
FOXA1 G275X enriches POU2F1 and GATA3 transcription factors to maintain luminal identity. (**A**) GATA3 and POU2F1 expression with non-targeting or sgRNA guides. (**B**) Expression levels of *Gata3* and *Pou2f1* in genotypes showing decreased expression of Gata3 in both knockouts. (**C**) UMAP with cluster labels of scRNA-seq expression data. (**D**) Barplot showing percentage of genotype within clusters. (**E**) 2D scatter plots of assigned basal and luminal scores for each genotype’s cells. Densities are calculated for each showing different enrichment per quadrant i.e. double positive, luminal, double negative, basal (I, II, III, IV). (**F**) ECDF shows probability of cell to be L1 vs cumulative proportion of cells per genotype. G275X contains the greatest number of cells likely to be L1, second most likely is *Gata3* KO, third is *Pou2f1* KO, and EV with lowest probability of L1 cells. (**G**) scATAC-seq UMAP with cell clusters labeled. (**H**) Pou2f1 ChIP and snATAC-seq tracks for Empty Vector-EV (purple) and GX275-GX (red) show two Pou2f1 ChIP-seq peaks within 10KB upstream of the Gata3 locus (shown in IGV). (**I**) Heatmap of chromVAR motif accessibility z-scores hierarchically clustered. Motifs were selected based on high accessibility and correlated expression per cell. (**J**) Umaps of chromatin accessibility (left) and corresponding gene expression (right) for Foxa1 (top) and Ehf (bottom). *Foxa1* expression increased in G275X *Pou2f1* KO cluster although accessibility is like other G275X clusters. The ETS transcription factor Ehf (bottom) has increased accessibility and expression in the Empty Vector clusters.

**Fig S5.**
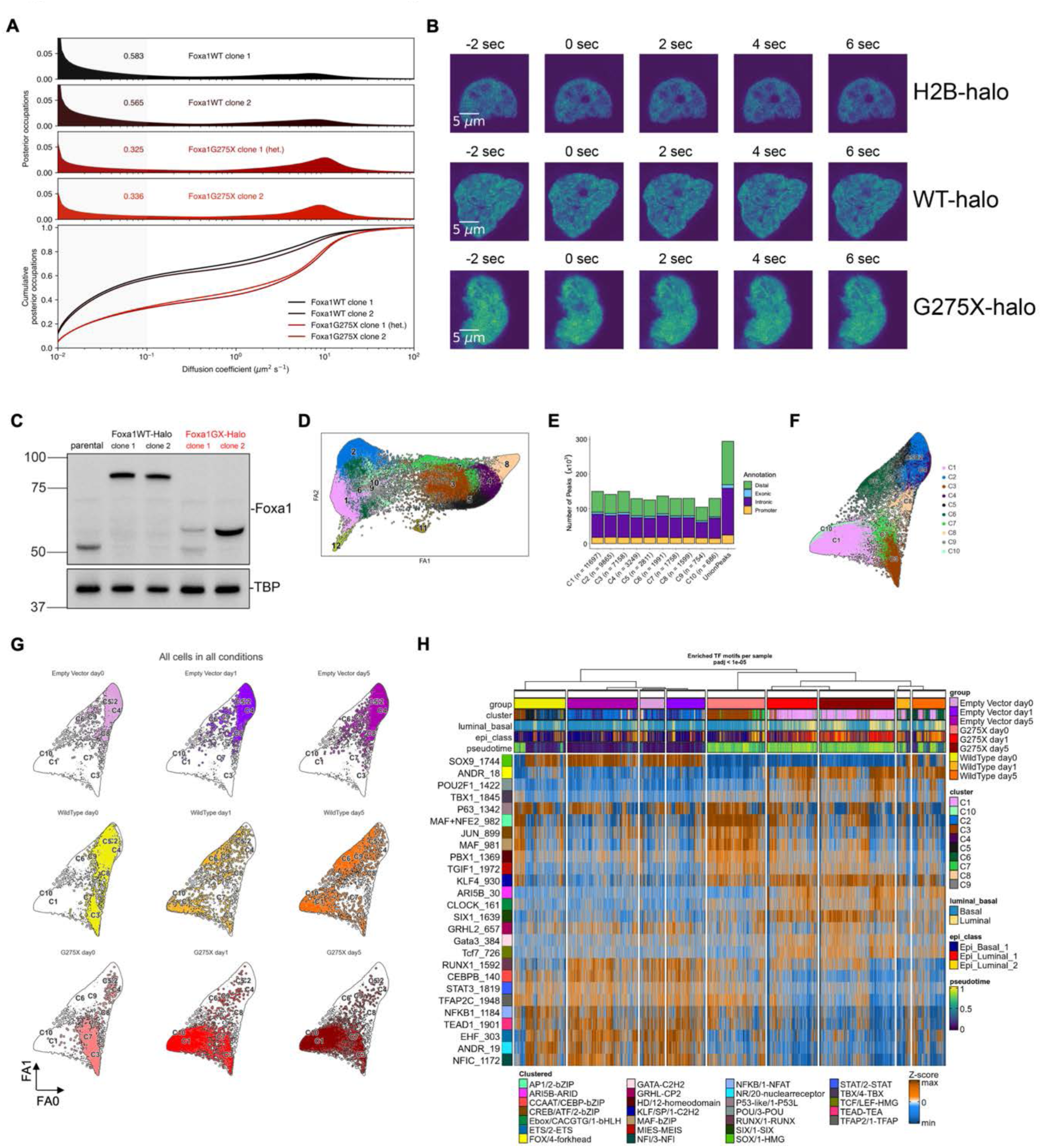
Faster activation of luminal identity in G275X mutant. (**A**) Experimental small particle tracking data (SPT) for WT and G275X with one replicate each. (**B**) Fluorescence recovery after photobleaching (FRAP) was performed on clonal LnCAP-AR lines, as well as the parental LnCAP-AR line exogenously expressing H2B-HaloTag as a control. (**C**) Western blot of parental, FOXA1 WT-halo and FOXA1 G275X-halo tagged proteins. (**D**) scRNA-seq force directed layout (FA) colored by leiden clustering. (**E**) Number of peaks within scATAC-seq clusters annotated with peak categories. (**F**) scATAC-seq FA with cells colored by cluster. (**G**) scATAC-seq FA per genotype and day. (**H**) scATAC-seq heatmap clustered by genotype and day. Top motifs are labeled on the left with chromVAR z-scores shown per cell (columns).

